# An implicit lipid model for efficient reaction-diffusion simulations of protein binding to surfaces of arbitrary topology

**DOI:** 10.1101/702845

**Authors:** Yiben Fu, Osman N. Yogurtcu, Ruchita Kothari, Gudrun Thorkelsdottir, Alexander J. Sodt, Margaret E. Johnson

## Abstract

Localization of proteins to a membrane is an essential step in a broad range of biological processes such as signaling, virion formation, and clathrin-mediated endocytosis. The strength and specificity of proteins binding to a membrane depend on the lipid composition. Single-particle reaction-diffusion methods offer a powerful tool for capturing lipid-specific binding to membrane surfaces by treating lipids explicitly as individual diffusible binding sites. However, modeling lipid particle populations is expensive. Here we present an algorithm for reversible binding of proteins to continuum surfaces with implicit lipids, providing dramatic speed-ups to many body simulations. Our algorithm can be readily integrated into most reaction-diffusion software packages. We characterize changes to kinetics that emerge from explicit versus implicit lipids as well as surface adsorption models, showing excellent agreement between our method and the full explicit lipid model. Compared to models of surface adsorption, which couple together binding affinity and lipid concentration, our implicit lipid model decouples them to provide more flexibility for controlling surface binding properties and lipid inhomogeneity, and thus reproducing binding kinetics and equilibria. Crucially, we demonstrate our method’s application to membranes of arbitrary curvature and topology, modeled via a subdivision limit surface, again showing excellent agreement with explicit lipid simulations. Unlike adsorption models, our method retains the ability to bind lipids after proteins are localized to the surface (through e.g. a protein-protein interaction), which can greatly increase stability of multi-protein complexes on the surface. Our method will enable efficient cell-scale simulations involving proteins localizing to realistic membrane models, which is a critical step for predictive modeling and quantification of *in vitro* and *in vivo* dynamics.

## I. Introduction

Proteins localize to membranes via multiple different binding modes, including recognition and binding to highly specific lipid head-groups (e.g. PI(4,5)P_2_) [1], electrostatically driven adherence to negatively charged membranes [2] (as performed by many BAR-domain proteins) [3, 4], and binding to membrane-inserted proteins (e.g. Ras) [5, 6]. Once proteins have localized to the surface, they can be further stabilized by interactions with additional lipids or transmembrane proteins, and these subsequent binding events are effectively two dimensional (2D) in their search [6, 7]. Capturing these various forms of membrane binding is critical for effective kinetic and spatial modeling of cell-scale systems with quantitative comparison to experiment, both in equilibrium and non-equilibrium systems. Because molecular modeling approaches that could capture the fundamental electrostatic or hydrophobic nature of these interactions are too expensive at this scale [8, 9], rate-based approaches such as reaction-diffusion provide a powerful alternative [10–14]. These modeling approaches can be used not only to study systems with known experimental binding constants [15], but as a means of fitting and extracting information from new experiments on protein-membrane interactions. We present a new algorithm for efficient rate-based and reversible binding of proteins to flat or curved continuum membranes. We then characterize how treating lipids as explicit particles versus an adsorptive surface can alter reaction kinetics and equilibria, and illustrate our implicit lipid model’s ability to recover behavior of the full explicit lipid method.

Our model offers a hybrid explicit-particle scheme that combines the efficiency of surface adsorption models with the enhanced spatial and molecular detail of single-particle binding sites. In a simple surface adsorption model, the surface is a boundary where adsorption occurs upon collision using an adsorption rate *κ* with units of length/time (see, e.g. Ref [16]). The more expensive but ultimately more detailed and flexible approach is to give the surface a density of reactive particles, and solution species can react according to a bimolecular reaction rate *k_on_* with units of volume/time. This second model, generally known as the Langmuir model [17], assumes a 1:1 binding stoichiometry between the surface species and the solution species. Both surface adsorption [16, 18–23] and Langmuir [7, 11, 23, 24] models have been used for single-particle reaction diffusion (RD) and for partial differential equation (PDE)-based modeling [25, 26]. The Langmuir model has a clear advantage in that it treats surface binding sites as a variable, accounting for heterogeneity and exclusion of other solution particles from binding occupied sites. This second model is also the only one that can naturally account for subsequent binding to surface sites as a 2D reaction, that is, after a solution particle has first been localized via a separate binding site [7]. However, for single-particle RD, propagating individual membrane binding sites (such as lipids or patches of lipids) is quite costly. Lipids are quite abundant. For example, at about 1% mol at the plasma membrane, the number density of PI(4,5)P_2_ is 25000/*μ*m^2^ [27], translating to about 50-80*μ*M given a yeast cell surface area and volume.

While more expensive than continuum PDE [25, 26] or stochastic RDME approaches [28–31], single-particle RD simulations are unique in capturing excluded volume[13, 14, 32, 33], capturing fluctuations driven by small copy numbers and diffusional collisions[11, 13, 14, 34], and enabling simulations of macromolecular self-assembly [35], thus providing a mechanism to bridge cell-scale modeling with molecular modeling approaches [36]. For single-particle binding to surfaces, our method provides critical advantages over existing adsorption methods. Several algorithms for surface adsorption based on discretizing the diffusion equation require short time-steps to provide accurate solutions [12, 20, 21, 37]. Methods using analytical solutions to the diffusion equation with a reactive boundary provide accurate solutions even for larger steps [16, 18, 19, 23, 38], but have restricted parameter regimes [16], are model specific [38], and in all cases, are derived only for planar surfaces[12, 16, 18–21, 23, 38] [37]. Surfaces with mixed boundaries (inhomogeneity) have been considered for receptor arrays on planar surfaces [38, 39], but these receptor disks are immobile. Perhaps most importantly, unlike these existing surface adsorption models, our implicit lipid model treats the density of the target lipid as a variable, thus naturally retaining the ability of the explicit lipid model to capture binding occupancy, non-homogeneous distributions, and 2D lipid binding. Critically, it applies on flat and curved surfaces, retains accuracy with large time-steps, and applies to reversible binding. This approach also provides a natural way to incorporate single-particle RD with hybrid continuum approaches[40–42], where the surface density is naturally a continuous variable. We achieve this versatility in our model by deriving reaction probabilities for a particle in solution to bind to a surface distribution of lipids that replaces individual binding sites with an approximation by a continuous field (Fig. 1). By defining reaction probabilities based on the Green’s function solution to the Smoluchowki model, we can take relatively large time-steps (~*μ*s) while correctly accounting for all possible collisions with the surface. Importantly, we propagate solution particles using simple Brownian updates, such that our method can be integrated with most single-particle simulation tools[11, 12, 14, 24]. The desorption probabilities we define enforce detailed balance to reproduce the proper equilibrium distribution for reversible reactions. Our approach applies to all types of surface models, with any reaction parameters, and can be widely adopted by existing single-particle methods.

**Figure 1.**
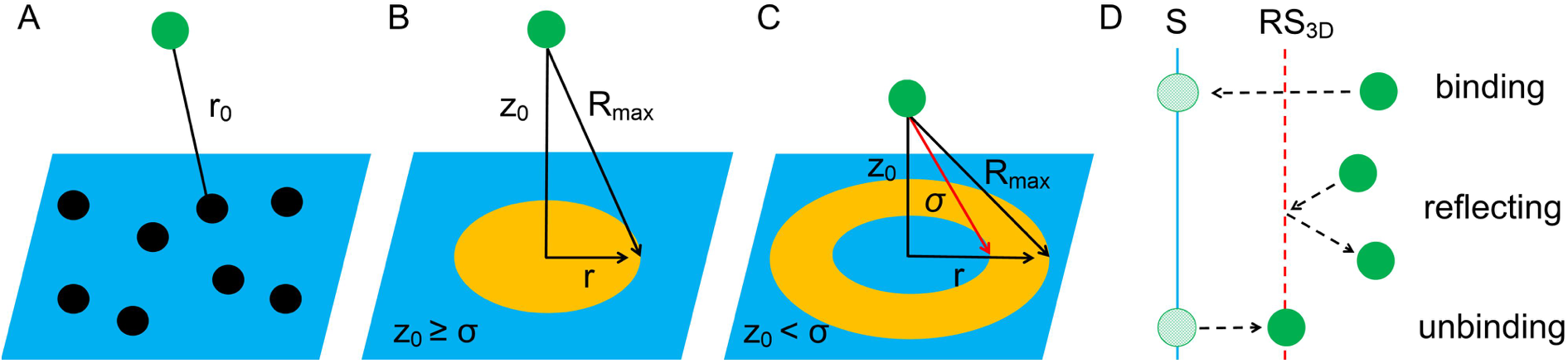
Schematic diagrams of the implicit lipid model. Binding sites or lipids on the surface, indicated by the *N_L_* black points (A), are replaced by a uniform field with density *ρ*_L_=*n*_L_/S (B and C). For a solution particle (green point), instead of evaluating binding to each of the lipid within a separation *r*_0_ < *R*_max_, we can instead evaluate the binding to the surface by integrating over all positions on the surface that are accessible (*r*_0_ < *R*_max_) given a height *z*_0_ (B). The separation 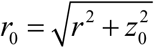 cannot fall below *σ*, hence a donut-shaped patch on (C). (D) Effects of the reflecting-surface *RS*_3*D*_. A diffusing particle cannot pass through the reflecting-surface, unless a reaction binding occurs in which case it is put on the membrane surface (denoted as S) directly. The dissociated particle is moved from the membrane surface and put on the reflecting surface.

By comparing adsorption and explicit lipid models of surface binding, we show how the choice of model can lead to significant differences in the kinetics of membrane association and occupancy at the membrane. Surface adsorption models can be designed to recover the same equilibrium occupancy of the Langmuir model simply by making the adsorption rate *κ* a function of lipid density, *κ*(*ρ_L_*) = *k_on_*·*ρ_L_*. This useful formula highlights the relationship between the two models, and guides comparison between them. As suggested by this formula and shown below, for parameter regimes with a high density of surface binding sites, both models produce nearly identical results. With limited binding sites, explicit lipid models produce substantially slower kinetics. Our implicit lipid method, on the other hand, shows excellent agreement with the kinetics of explicit lipid models. We further compare the microscopic explicit lipid model with macroscopic binding models using spatially unresolved ordinary differential equations (ODEs). In small and rate-limited systems, these macroscopic rate equations produce quite comparable kinetics to the microscopic kinetics of the explicit lipid model, with results diverging for diffusion limited reactions.

By establishing kinetic data for a range of reaction parameters and surface topologies, from simple (planar) to complex using spatially resolved methods, we provide a standard for the comparison of other rate-based models across simulation software implementations. Curved surfaces present several challenges relative to flat surfaces, particularly if they are not analytically defined (e.g. a sphere or cylinder). We achieve binding to surfaces of arbitrary topology by using a sublimit surface mesh model [43], and by defining accurate methods for detection of the surface of and reflection off the surface [44, 45], and diffusion on the surface [46]. In this paper, we first introduce the single-particle model for binding to explicit particles embedded in surfaces, before deriving our new implicit lipid model and the associated algorithm. We define relationships between rates of surface adsorption models, explicit lipid and our implicit lipid model to establish accurate and comparable definitions of macroscopic rates. We show how the models produce distinct kinetics in low lipid regimes, with converging kinetics at high surface coverage. Our method can be incorporated with both existing single-particle methods and hybrid PDE based models, as the solution particles can bind to a spatially dependent density of surface particles. We further demonstrate how 2D binding to surface sites (i.e. via localization by separate interfaces), can be done with our implicit lipid model, which is not possible with surface adsorption models. We apply this method to heterogeneous densities of surface lipids, and to curved mesh surface models as an important step in integrating RD methods with continuum deformable membrane models [43]. Finally, we note that while none of these rate-based models capture molecular properties of lipid bilayers (e.g. electrostatic attraction and repulsion), some molecular features are implicitly represented by the lipid density and binding rate. This simpler model is a necessary starting point for ultimately capturing how binding responds to collective changes of the surface, such as curvature [3, 47, 48].

## II. Theory

### A. Single-particle reaction-diffusion: solution particles binding to explicit lipids on surfaces

#### A1. Binding to lipids on a flat surface

Binding between a particle in solution at ***r***_1_ to a lipid particle restricted to a reflective 2D surface at ***r***_2_, thus transitioning the 3D solution particle to the 2D surface, follows the same general theoretical approach as between two particles in solution (Fig 2). The Smoluchowski model for binding between reactive pairs has been quite effectively used for single-particle RD with large time-steps [13, 14, 24]; we define the model here as there is a modification required relative to standard treatments of two solution particles [49, 50], due to the reflective surface. The diffusion equation for the particle pair can be written as

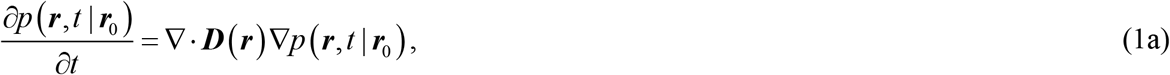

with their separation ***r*** = ***r***_1_ − ***r***_2_ and initial separation ***r***_0_, and *p*(***r***,*t*|***r***_0_) is referred to as the Green’s function, (GF). The diffusion tensor ***D***(***r***)=***D***_1_+***D***_2_ is a generally non-isotropic matrix because the diffusion coefficients on the surface are non-zero in only 2 dimensions (for 2D diffusion on a horizontal plane, *D*_2z_ = 0). The reaction is modeled via the reactive boundary at *r* = *σ*,

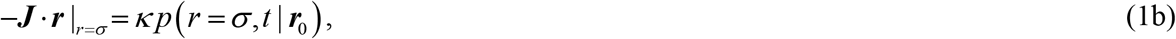

where ***J*** = −***D***(***r***)*p*(***r***_12_,t|***r***_120_) is the flux at ***r***, and ***r*** is the direction vector of ***r***. The reactivity *κ* occurs over the accessible surface of a sphere of radius *σ*, such that *κ* = *k_a_*/*B*(*σ, d*), where *k*_a_ is the microscopic (or intrinsic) binding rate constant. Unlike in a full three dimensional (3D) solution, where *B*(*σ*,*d* = 3) = 4*πσ*^2^, here only a hemi-sphere is accessible, 2*πσ*^2^, due to the additional boundary of the reflective surface,

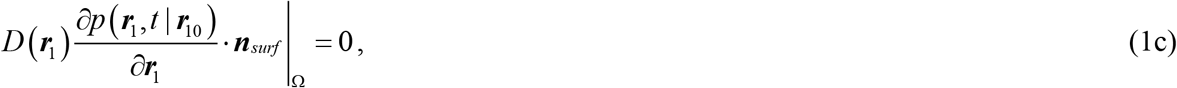

where Ω indicates the boundary surface, and ***n**_surf_* is the direction vector of surface. This affects the macroscopic rate as described in Section C below. The other boundary condition is

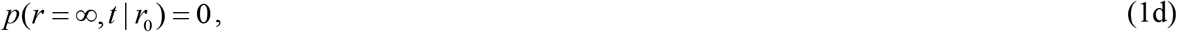

and the initial condition is a delta function: *p*(*r, t* = 0| *r*_0_) = *δ*(*r* − *r*_0_)/*B*(*σ,d*).

**Figure 2.**
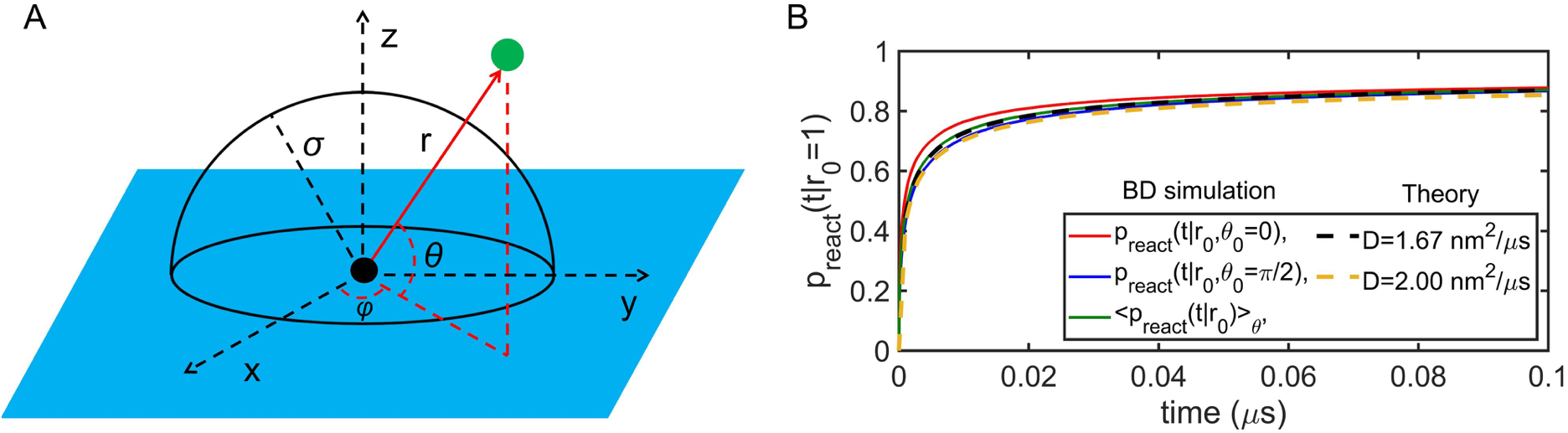
Model for binding to explicit lipids on a flat surface. (A) Geometric schematic in the spherical coordinate system. The green point represents a protein particle in solution, and the black point represents the lipid particle on membrane. The green particles reflect off the membrane surface (blue color). This constrains the polar angle *θ* to be between 0 and *π*/2, while the radius *r* and azimuth angle *φ* are not influenced. (B) Reaction probability between the particles depicted in (A) as a function of time, given an initial separation *r*_0_=1nm. Results from Brownian Dynamics simulations of (A) are shown in red and blue, at different initial polar angles, with the average over all orientations in green. The theoretical solution provided by Eq. 2 is shown in black dashed, agreeing extremely well with the BD average in green, using the weighted 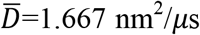. For comparison, we show the theory with *D*=*D*_1_x+*D*_2_x=2 nm^2^/*μ*s, which is slightly lower than the average. Results for *k_a_*=347.18 nm^3^/μs, *σ* = 1 nm.

For a lipid particle diffusing in 2D in a planar surface, the problem is perfectly symmetric around the plane, e.g. at *z* = 0. Reflection simply flips the polar angle *θ* to remain between 0 and π/2 without affecting *r* or *φ*, (Fig 2a). The Green’s function (GF), *p*(***r***,*t*|***r***_0_) can thus be defined from the full 3D problem, with the only complication being the anisotropy of the diffusion matrix ***D*** (*D_x_*=*D_y_*≠*D_z_*). Unfortunately, although the diffusion anisotropy can be removed by a change of variables, this then introduces an anisotropy on the radiative boundary condition (the *z*-axis is rescaled) [23], and thus the GF can only be solved exactly as a function of initial and final particle separation *and* angular orientation.

Instead, we will approximate the dynamics as occurring under an isotropic 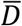, weighted by the *x*, *y*, and *z* components, so that our algorithms can use the analytically known GF solution in 3D dependent only on particle separation *r*, *p*(*r*,*t*|*r*_0_) [14]. As usual, the reaction probabilities for a pair of particles at a separation of *r*_0_ are defined based on the survival probability, such that

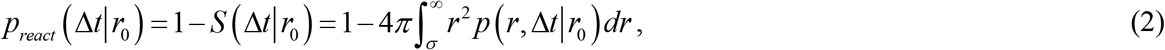

producing the same functions used in 3D. To define 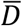, we note that the anisotropic diffusion in *x, y*, and *z* means that the diffusion in spherical coordinates along the relative separation *r* is a function of the angular orientation, *D*(*θ, φ*) = *D_x_*cos^2^(*φ*)sin^2^(*θ*)+*D_y_*sin^2^(*φ*)sin^2^(*θ*)+*D_z_*cos^2^(*θ*). Since *D_x_* = *D_y_*, then we have *D*=*D_x_*sin^2^(*θ*)+*D_z_*cos^2^(*θ*). If we weight *D* over all polar angles, then 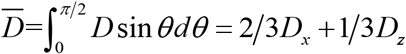. We verify that using the known 3D reaction probabilities with this D provides excellent agreement with the exact reaction probabilities determined using Brownian Dynamics simulations (Fig 2b). Brownian Dynamics simulations were run using the method of Zhou [51, 52], similar to recent work [35, 53], with the addition of the reflecting surface around the lipid particle.

#### A2. Binding to explicit lipids on a curved surface

For binding to a lipid diffusing on a curved surface, the system is no longer perfectly symmetric across the reflecting surface, and thus the reaction probabilities are not exactly described via the GFs discussed above. Here, we will simply approximate the reaction probabilities based on the GFs described above, thus ignoring the effect of the local curvature on the reactive flux at *r* = *σ*. The curvature of the reflecting surface would have two main effects on the GF solutions. First, reflection off a concave surface, for instance, tends to redirect particles closer to the surface compared to reflections off flat or convex surfaces, resulting in a small enhancement to reactive collisions. Second, for a concave surface, the accessible surface of a sphere of radius *σ* around the lipid particle would be slightly less than the hemisphere for a flat surface, resulting in a small reduction in net reactive flux. These effects are thus somewhat compensatory. An exact GF and reaction probability would depend on the full orientation between particles, and the local geometry of the reflective surface, requiring a costly numerical solution. Instead, the effect of the curvature is reduced by propagating a smaller time-step, rendering the flat-surface GF solution above a more and more accurate approximation. In our results below, we find that evaluating binding probabilities to a particle on a curved surface assuming a flat reflection model of Section A1 does not introduce statistically measurable deviations in the known equilibria, suggesting it is a good approximation.

#### A3. Dissociation

As performed in previous work [35], dissociation between explicit particles is treated as a Poisson process,

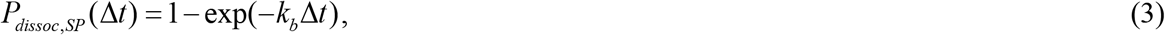

where *k_b_* is the microscopic dissociation constant. To recover detailed balance, and thus the proper equilibrium, dissociation must return particles to the same separation where the reaction occurs, which for the Smoluchowski model is at contact, *r* = *σ*.

### B. Implicit lipid model: explicit solution particles binding to implicit lipids on surfaces

The explicit lipid model for binding to surfaces described above achieves high resolution of the spatial distribution and its dependence on diffusion. However, it is expensive to propagate all of the lipid particles; as small, densely packed constituents of membranes, they are often much more numerous than solution molecules. Our new implicit lipid model replaces the lipid particles by a field or density distribution (see Fig. 1). This approach reduces the spatial detail of the lipids, but eliminates the need to propagate the lipids on the membrane surface. Diffusion of solution particles can still be propagated using simple Brownian updates, and binding probabilities are closed-form analytic functions for planes and spheres. An important aspect of this approach is that the depletion of the lipid particles due to binding of proteins can be captured in the lipid density, *ρ*_L_ = *n*_L_/*S* (*n*_L_, the number of unbound lipids; *S*, the area of membrane surface). Once a binding event occurs, *n*_L_ decreases, which will reduce the density and change the reaction probability for subsequent events. Dissociation events return lipids to the pool and increase the density. By integrating over a surface patch of the membrane accessible to the solution particle, thus considering all possible collisions with the surface, this model can be applied to surfaces of varying curvature, lipid density, and to particles restricted to 2D. We enforce detailed balance by specifying an additional reflection surface parameter, *RS*.

#### B1. Derivation of the implicit lipid model for a planar surface

For a solution particle that interacts with a field of lipid particles located on a planar surface at *z* = 0, where first we assume a fixed density of *ρ*_L_=*N*_L_/*S*, we can define the implicit lipid survival probability for the solution particle in a time Δ*t*, given a height above the membrane of *z*_0_, by integrating over all collisions with the surface (Fig 1). This is given by the product of the pairwise survival probability between the solution particle and each position on the surface, weighted by the lipid density:

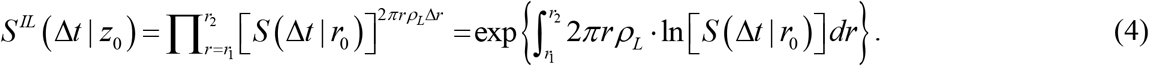

where, from the geometry of the problem, we define 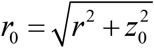, and *r*_1_ = 0 if *z*_0_ ≥ *σ*, 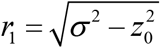 if *z*_0_ < *σ* (see Fig. 1). The well-known reaction probability *p_react_* (Δ*t*|*r*_0_)=1−*S*(Δ*t*|*r*_0_) (Eq. 2) can only be evaluated for *r*_0_ ≥ *σ* [14] (Fig 2c). The upper integration limit *r*_2_ can be set to infinity, although it can be convenient to truncate the integral. Because the integrand drops to zero rapidly, we define a cutoff based on the 3D distance where the particles are too far apart to collide in a time-step, 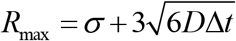, which gives 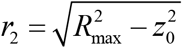, where 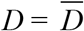 (section A1).

Then the reaction probability for the implicit lipid model to a planar surface is written as:

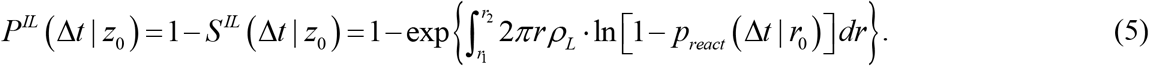

This integral must be solved numerically. However, if the time step Δ*t* is relatively small (order of *μ*s) and the lipid density *ρ*_L_ on the membrane is relatively low, then the reaction probability is very well approximated analytically as:

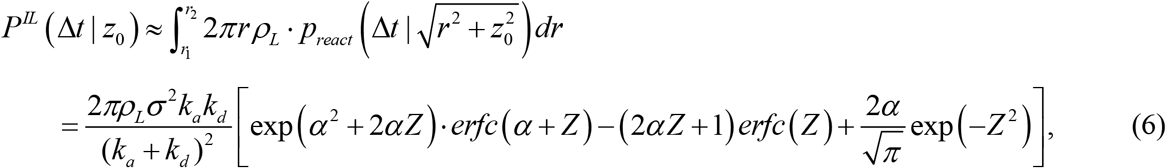

where 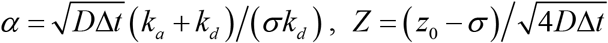, *k_d_* = 4π*σD*, and *k_a_* is the intrinsic binding rate. When *z*_0_ < *σ*, the probability is independent of the specific value of *z*_0_, and is in fact always given by *P^IL^*(Δ*t*|*z*_0_=*σ*). A typical curve of *P^IL^*(Δ*t*|*z*_0_) is shown in Fig. 3, where Eq 6 is indistinguishable from the numerical solution to Eq 5, and is faster to evaluate. This approximation works because one can show that Eq 5 is given by Eq 6, minus corrections to prevent multiple events per step. These corrections are negligible for small Δ*t*.

**Figure 3.**
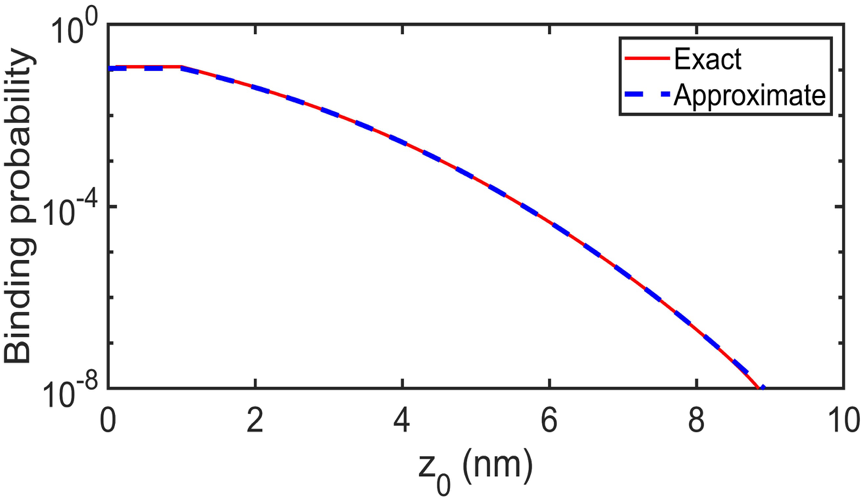
The reaction probability for the implicit lipid model, *P^IL^* (Δ*t*|*z*_0_) (red--Eq. 5) can be extremely well approximated by a closed form analytical function (blue dashed-Eq. 6). Parameters: *σ* = 1 nm, 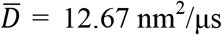, Δ*t* = 0.1 μs, *ρ_L_* = 0.025 nm^−2^, *k_a_* = 347.18 nm^3^/μs and *k_b_* = 2.09 s^−1^.

The dissociation of particles from lipids follows a Poisson process,

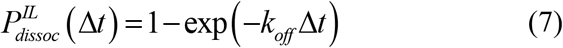

where unlike the explicit lipid model, here we must use *k_off_*, the macroscopic dissociation rate that satisfies *k_off_* = *k_on_K_D_* = *k_on_* · 2*k_b_*/*k_a_*, and *k_on_* is a simple function of *k_a_, D*, and *σ* as described in Section C below. This is necessary because the explicit lipid method returns particles exactly to contact, where the possibility of geminate rebinding creates a distinction between *k_b_*, the intrinsic off rate, and *k_off_*, the macroscopic off rate. For rate-limited reactions *k_b_* and *k_off_* are the same, but for diffusion-influenced reactions where *k_a_* is large, *k_b_* is faster than *k_off_*(*k_b_* ≥ *k_off_*). Here, there are no explicit lipids to drive geminate rebinding, and thus the dissociation follows the macroscopic rate.

The explicit solution particles are propagated using free diffusion. Unlike in the FPR algorithm [14], here we do not reweight the reaction probabilities, as the position updates for particles that did not bind result from an effectively many-body interaction with the surface, rather than a pairwise reaction. To guarantee the correct equilibrium in the implicit lipid model, one final parameter is needed, the position of the reflecting surface, which excludes particles from getting within a distance *RS*_3D_ of the surface. Detailed balance requires that at the equilibrium state, during a time step Δ*t*, the number of solution particles binding to the surface should be equal to the number of particles dissociating from the surface. This is expressed as:

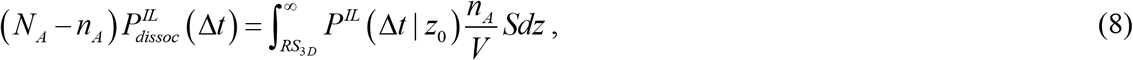

where *V* is the volume of system, *S* is the area of membrane surface, *N*_A_ is the total number of solution particles, *n*_A_ is the number of unbound solution particles at equilibrium state, and *RS*_3D_ is the position of reflecting surface above the membrane. The above function has an analytical solution:

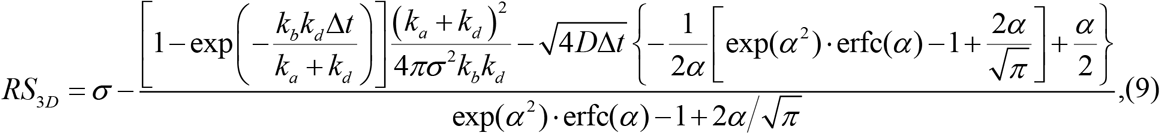

where *α* and *k_d_* are same as in Eq. (6). If both *k_b_* and Δ*t* are relatively small, then *RS*_3D_ has the simple expression:

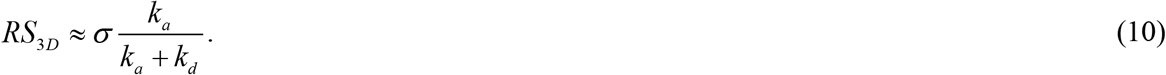

where *k_d_* = 4*πσD*. We can see that *RS*_3D_ has a value of 0 < *RS*_3D_ < *σ*. The reflecting surface depends only on *σ*, *k_a_* and diffusion constant *D*. It does not depend on the system size or particle numbers, as we would expect for this reaction model.

In sum of the implicit lipid model, solution particles move according to free diffusion and measure distances to the membrane surface proper, but their position updates must exclude them from occupying the layer *RS*_3D_ above the membrane, via reflection. When dissociation occurs, particles are returned to a distance of *RS*_3D_ above the membrane surface. Then, using the reaction probability of Eq. 6 and the dissociation probability of Eq. 7, we recover the correct equilibrium and kinetics very similar to the explicit lipid model, as we show below.

#### B2. Implicit lipid model: modification for a curved membrane

If the membrane is not planar, the binding probability must account for the curvature of the membrane. This alters the distance relative to the rule used above, where previously 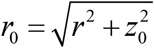. Now, we use the more general expression for Eq (6):

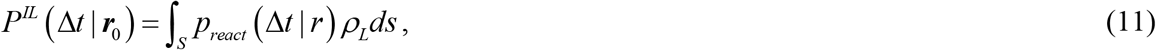

where *S* represents a patch on the surface accessible to the diffusing solution particle within a step Δ*t*, ***r***_0_ represents the spatial position of the particle in solution, and *r* is the distance from the particle to each point within the surface patch. Here again, *S* need not be the entire surface, as the reaction probability will drop to zero for *r* > *R_max_*. Also, the surface integral must be evaluated over distances *r* ≥ *σ*. This integral must be computed numerically to account for the changes in distances over the surface patch, but can be thought of as the average of the reaction probabilities over all separations, multiplied by the lipid density and the surface area of the patch. For a spherical surface, the reaction probability in Eq.11 is analytically solvable, providing faster evaluations of binding probabilities (Supplemental Information).

The dissociation probability is not dependent on curvature. However, the reflecting surface *RS*_3D_ depends on the reaction probability, Eq. (11), which changes slightly with curvature. Following the conservative detailed balance, we need to combine Eqs. (7), (8) and (11) to calculate the value of *RS*_3D_. In the SI, we show the calculation for binding to both the interior and exterior of a sphere (positive and negative curvature), illustrating that *RS*_3D_ can be reproducibly simplified to the same result of Eq. 10. One can thus generally assume that when the membrane curvature is not extreme over the length of the time-step, Eq. (10) is a very good approximation of *RS*_3D_.

#### B3. Implicit lipid model for 2D reactions

An important feature that cannot be captured by surface adsorption models is binding to lipids by proteins that are already localized to the surface. This can happen either because proteins are localized to the surface through a protein-protein interaction, or because a protein has multiple lipid binding sites. For these 2D reactions, the relative displacement *z*_0_ above the membrane is always zero, such that, using the simpler formulation of Eq 6, the integral is given by:

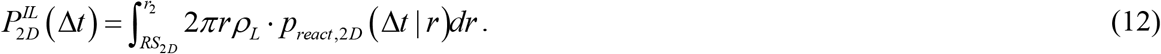

Here again the upper limit of integration can be chosen to a point where the reaction probability drops to zero, at 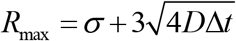. *RS*_2D_ is the reflecting distance (we will introduce it below), which similar to 3D, is a parameter used to enforce detailed balance. The reaction probability requires using the known solution to the 2D association problem, *p*_*react*,2D_(Δ*t*|*r*), (see [53]) which is itself an integral over Bessel functions. By taking *p*_*react*,2D_(Δ*t*|*r*) into Eq. (12), the binding probability is expressed as:

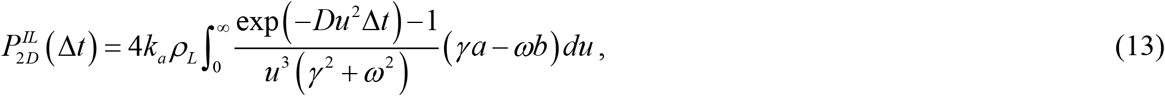

where *γ* = *huY*_1_(*uσ*)+ *k_a_Y*_0_(*uσ*), *ω*=*huJ*_1_(*uσ*)+*k_a_J*_0_(*uσ*), *a* = *uR*_max_*J*_1_(*uR*_max_) − *uR*_2*D*_ *J*_1_(*uRS*_2*D*_), and *b* = *uR*_max_*Y*_1_(*uR*_max_) − *uR*_2*D*_*Y*_1_(*uRS*_2*D*_), *h*=2*πσD*. *J*_n_ denotes order *n* Bessel function of the first kind, and *Y*_n_ denotes order *n* Bessel function of the second kind. The integral must be solved numerically. We use the GNU scientific library (GSL) quadrature routines for numerical evaluation.

The dissociation of particles from lipids still follows a Poisson process,

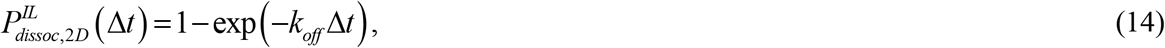

where *k_off_* is the macroscopic dissociation rate that satisfies *k_off_* = *k_on_K_D_* = *k_on_k_b_*/*k_a_*, and a macroscopic association rate *k_on_* for 2D is determined based on our previous work [53],

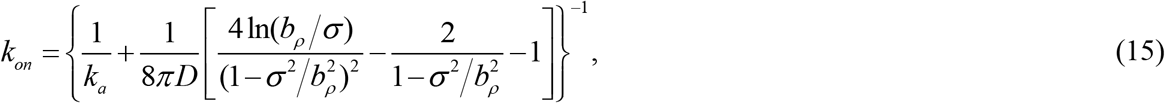

Where 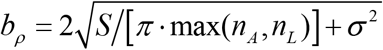, and *S* is the membrane area, *n_A_* is the number of unbound particles, *n_L_* is the number of unbound lipids.

Similar to the 3D case, we need to introduce a reflecting distance, *RS*_2D_, that will enforce detailed balance between binding and unbinding reactions, which will be ≥ *σ*. At the equilibrium state, during a time step Δ*t*, the number of particles binding to lipids must be equal to the number of particles dissociating from lipids, which can be expressed as:

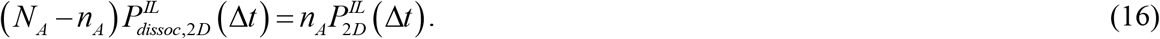

Taking Eqs. 13-15 into the equation above, and considering that *k_b_* is usually small, we have

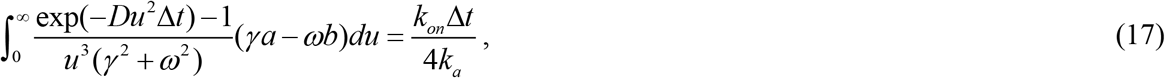

where *γ, ω, a, b* are defined as before in Eq. 13 (*a* and *b* are functions of *RS*_2D_). By numerically solving Eq. 17, we obtain the value of *RS*_2D_. It is usually larger than *σ* (Fig. S1).

On a curved surface the approach is similar to the 3D case, where we explicitly integrate over all points along the surface in terms of their geodesic distance from the protein particle:

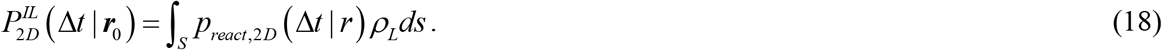

Note that the value of *r* will start from *RS*_2D_. The dissociation is still defined by Eq. 14, and the reflecting distance *RS*_2D_ will be calculated by Eqs. 14, 16 and 18.

#### B4. Implicit lipid model for non-uniform lipid densities

In the above examples, the lipid density was used as a fixed variable *ρ_L_*. Non-uniform lipid densities could arise due to phase separation [54] or some other confining mechanism. Our method can straightforwardly accommodate such inhomogeneous lipid densities: the same integrals of Eqs. 5, 6, 11, or 18 apply but the integration over reaction probabilities is weighted by a spatially varying lipid density, *ρ_L_*(*r, φ*), where *φ* is the azimuthal angle. While this requires numerical integration even for Eq. 6, this enables application of this model to a hybrid PDE-based method with explicit solution particles, where the lipid density on the surface could be defined as a continuous and diffusive distribution. To facilitate direct comparison with the explicit lipid simulations, we demonstrate this approach where the lipid densities on the surface are restricted to zero in one region, and a fixed value *ρ_L_* in the other region.

### C. Macroscopic Rates from the microscopic Smoluchowski model

For both the explicit lipid model and the implicit lipid model that is derived from it, the relationship between the microscopic or intrinsic binding rate (*k_a_*) and the macroscopic rate *k_on_* is different than the known relationship for both particles in solution, due to the reflective surface. The total reactive flux across the radiation boundary at *r* = *σ* is reduced, and for a planar surface the reduction is by exactly 1/2. As a result, the macroscopic rate differs from the rate observed in 3D solution by a factor of 1/2 for the same value of *k_a_*. Specifically, the macroscopic rate is known to be related to the survival probability through [51]

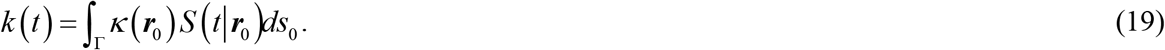

where ***r***_0_ are all the points on the reactive surface Γ. For a spherically symmetric interaction, the reactive surface is at *r*_0_ = *σ*, and *φ* = [0, 2π], with the reflective surface restricting the polar range to *θ* = [0, arccos(*σc*/2)], thus the total reactive surface area is 2*πσ*^2^(1 − *σc*/2). Here *c* represents the local curvature of the surface, which can be positive or negative (*c*=1/*R* for a sphere of radius *R*), or for a planar surface, *c* = 0. The survival probability is provided by the full 3D problem, where *κ*=*k_a_*/4*πσ*^2^. As a result,

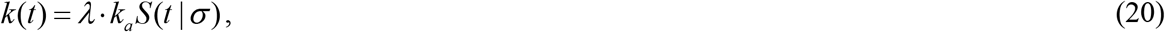

where the reduction factor *λ*=1/2 (1−*σc*/2). Given the well-known survival probability in 3D [14], the macroscopic rate is

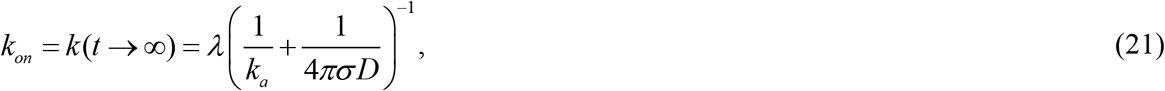

or *λ* times the rate that is produced in solution (1/2 for a planar surface). This means that for a single-particle RD simulation to produce a specific macroscopic rate, Eq. (21) must be used to extract the *k_a_* value, which will be up to a factor of *λ*^−1^ higher than for a solution reaction.

Critically, this results in a similar modification to define the equilibrium constant from the intrinsic rates, which applies for any reaction where one reactant is localized to a reflective surface. In 3D, we have that 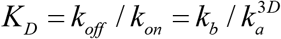. For transitions between solution and the surface, we have that

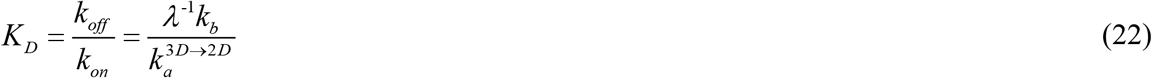

This requires care when reactant pairs both start off in solution with a specified 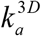, and one becomes localized to the surface. Subsequent binding reactions using 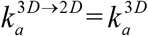 will follow the weaker *K_D_* of Eq. 22, despite involving the same two reactants. We noted in a recent publication that this correction is important in models of membrane localization to producing a proper equilibrium, rather than non-equilibrium, steady-state [7]. To account for this, we use a simple approach in our software. All *k_a_* values are defined based on a 3D reaction, and if one reactant is localized to the membrane, we multiply *k_a_* by the zero-curvature value of *λ*^−1^=2 to preserve the same value of 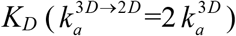, although this results in slightly different *k_on_* values. In Table 1 we list the relationships between the binding rates, unbinding rates, and equilibrium constant, for 3D, 2D and the 3D→2D transition. In Fig. S2 we illustrate the application of these relationships by comparing the microscopic explicit lipid simulations with an ODE solution using the macroscopic rates. Lastly, we note that the macroscopic rates for a curved surface will not be exactly given by these formula, as the survival probability and the size of the reactive surface change slightly with curvature.

**Table 1.** Relationships between microscopic and macroscopic reaction rates

| Systems | $3D$ | $3D \rightarrow 2D$ | $2D$ |
| --- | --- | --- | --- |
| Relation to 3D microscopic binding rate | $k_a^{3D}$ | $k_a^{3D \rightarrow 2D} = 2k_a^{3D}$ | $k_a^{2D} = k_a^{3D} / (2\sigma)$ |
| Microscopic dissociation rate | $k_b$ | $k_b$ | $k_b$ |
| Macroscopic binding rate | $k_{on} = \left( \frac{1}{k_a^{3D}} + \frac{1}{4\pi\sigma D} \right)^{-1}$ | $k_{on} = \frac{1}{2} \left( \frac{1}{k_a^{3D \rightarrow 2D}} + \frac{1}{4\pi\sigma D} \right)^{-1}$ | $k_{on} = \left\{ \frac{1}{k_a^{2D}} + \frac{1}{8\pi D} \times \left[ \frac{4\ln(b_\rho/\sigma)}{(1-\sigma^2/b_\rho^2)^2} - \frac{2}{1-\sigma^2/b_\rho^2} - 1 \right] \right\}^{-1}$ |
| Macroscopic dissociation rate | $k_{off} = k_{on} k_b / k_a^{3D}$ | $k_{off} = 2k_{on} k_b / k_a^{3D \rightarrow 2D}$ | $k_{off} = k_{on} k_b / k_a^{2D}$ |
| Equilibrium state | $K_D = k_b / k_a^{3D}$ | $K_D = 2k_b / k_a^{3D \rightarrow 2D}$ | $K_D = k_b / k_a^{2D}$ |

### D. Background on Surface Adsorption Models

In adsorption models, collisions with the surface result in ‘binding’ with an adsorption rate *κ* with units of length/time. To compare against models of bimolecular association to surface sites with rate *k_on_* (the explicit or implicit lipid models), we can use *κ*=*ρ_L_k_on_*, with *ρ_L_* being the density of lipids per area. We show here results both where *κ* is fixed, and where *κ*(*ρ*_L_(*t*)) varies as the lipid density changes in time. Desorption is treated like a standard dissociation reaction. An efficient and spatially dependent model for adsorption/desorption in single-particle reaction-diffusion was introduced by Andrews in Ref. [16]. Because this method cannot be used to simulate reversible adsorption when *κ* is low (requires *κ*^2^ ≥ 4*k_off_D*), it cannot be used for arbitrary binding strengths, as we do in some simulations here. We thus briefly describe the application here of the Smoluchowski model in 1D to surface adsorption, which has been already used in published algorithms [18, 23, 38].

#### D1. Adsorption using the Smoluchowski model in one dimension

For adsorption of a diffusing particle to a planar surface, the height above the surface is the only variable influencing the reaction. We can therefore describe the particle position based on the product of the free diffusion propagator in *x* and *y*, and diffusion in *z* subject to a radiation boundary at the surface, (e.g. *z* = *σ*). Specifically,

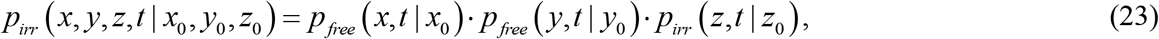

where 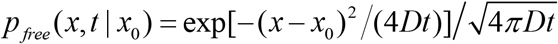, and same for *y*. The solution to the 1D Smoluchowki model *p*(*z,t*|*z*_0_), parameterized by the diffusion constant *D* = *D*_z_ and the rate *κ* (with adsorption units of length/time), is known to be [19, 49],

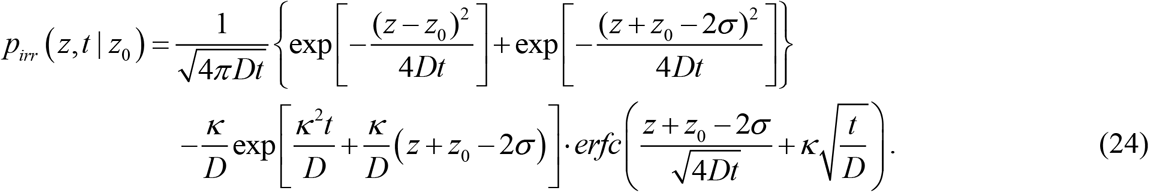

where *σ* can be used to move the surface to any arbitrary height. The reaction probability is then given by,

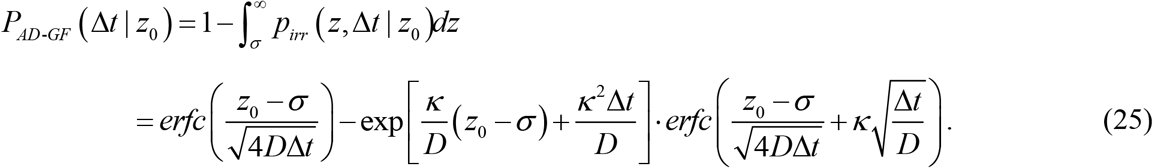

This probability depends on *z*_0_, which can then be compared with the result for *P*^IL^(Δ*t*|*z*_0_) derived above from the full 3D GF. The probability distributions are not identical (Fig. S3), which is not surprising, considering that one has 3D diffusion and the other does not. If we set *σ* = 0 for this model, for a small intrinsic binding rate *k_a_* they are comparable (Fig. S3a), but for a large *k_a_* they are more distinct (Fig. S3b). Therefore, it is perhaps not surprising that the many-body simulations we show below produce better agreement for rate-limited (small *k_a_*) reactions. Importantly, for a curved surface, the height above the surface is not independent of *x* and *y* diffusion, and thus this model would have to be adjusted to correct for this. We thus apply it here only to planar surfaces. Finally, for this single-particle method, desorption is again treated as a Poisson process, and the equilibrium for a solution population of A particles will be defined by *κ*/*k_desorb_* = [A]_*mem*_/ [A]_*sol*_.

## III. Methods

### A. Systems

Here, we set up four systems to represent different membrane surfaces or different characteristics of lipid distributions. Fig. 4a shows a planar surface in a cube, while in Fig. 4b the lipids are distributed non-uniformly, occupying only half of the bottom plane. Figs. 4c and 4d represent systems with curved membrane surfaces, both a sphere (4c) and a complex surface with four joined spheres that presents a range of curvature over convex and concave regions.

**Figure 4.**
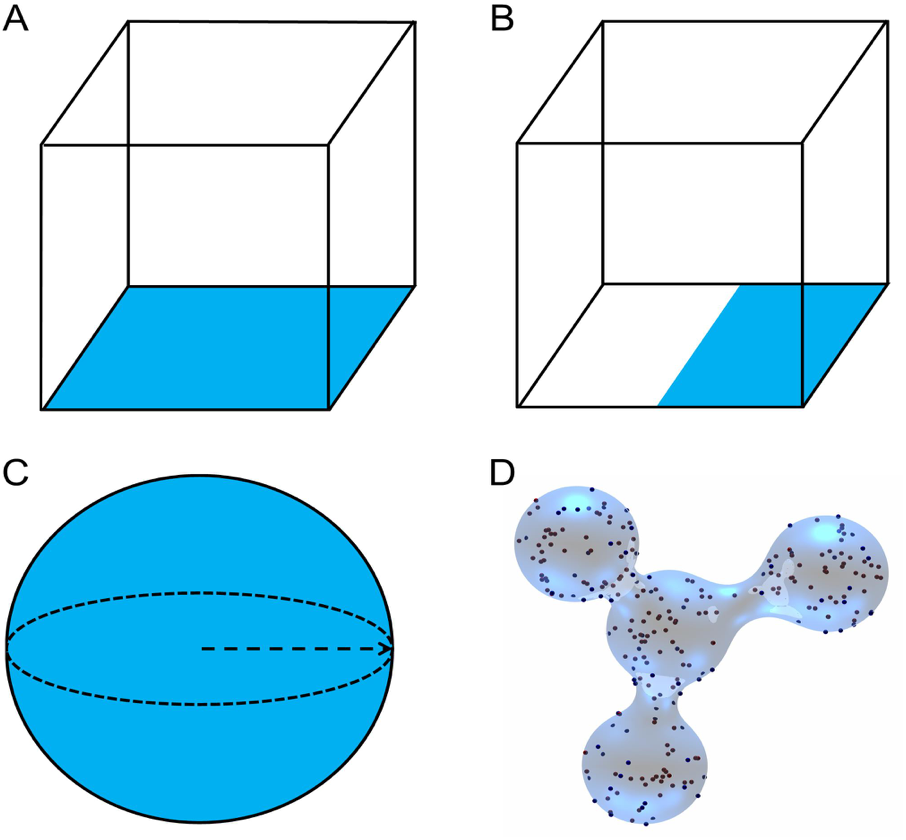
Schematics of four systems. A flat membrane surface (A), half-planar surface (B), spherical surface (C), and irregular curved surface (D) are shown. All the membrane surfaces are in blue color.

### B. Simulation details of the explicit lipid method

Simulations of the explicit lipid model in 3D and 2D were implemented here with the FPR algorithm [14, 35, 53]. Initial configurations were produced by a randomly sampled uniform distribution of particles in solution or on the surface. The binding probability of a solution particle to a lipid particle at a distance *r* < *R_max_* is determined by the pairwise association probability *p_react_*(Δ*t*|*r*) (Eq 2) multiplied with time-dependent reweighting ratios determined from the 3D GF [14]. In 2D, the same procedure applies, with the 2D GFs used. A uniformly distributed random number between 0 and 1 is generated and compared to the binding probability. If the random number is smaller than the binding probability, then the binding reaction occurs, otherwise the particle survives from this reaction and needs to check whether to bind to other lipid particles. If the binding takes place, the particle is put adjacent to the lipid particle on the membrane surface at a separation of *σ*. If a particle does not bind, then it is propagated according to the free diffusion, where its new position is rejected and resampled if the particle overlaps (*r* < *σ*) any reaction partners. Particles that cross the membrane or boundaries are reflected back inside the box. For bound species, the dissociation probability is evaluated by Eq. (3). If dissociation occurs, the particle becomes free and is left at contact with its former partner, at *r* = *σ*. Only in the next time step is it allowed to attempt a new reaction or diffuse.

### C. Simulation details of the implicit lipid model

For implicit lipid simulations with a flat membrane surface, if a solution particle is within *R*_max_ of the surface, based on the shortest distance to the surface, then the probability of binding will be evaluated by Eq. (6). Also, a uniformly distributed random number between 0-1 is generated and compared to the binding probability. If the random number is smaller than the binding probability, then the binding reaction occurs, otherwise the particle survives. If the reaction occurs, then the solution particle is placed on the membrane surface at the shortest distance. If the particle survives from binding, then it is freely diffused, where now the reflection occurs across the plane located at *RS*_3D_ above the membrane. For bound species, the probability of dissociation is calculated by Eq. (7). If the dissociation occurs, then the particle will be put at the closest point on the reflecting surface and will be treated as a free particle in the next time step. Once a binding or unbinding event occurs, the number of free lipids is updated, such that the density *ρ*_L_(*t*) of free lipids is correct at all times.

For 3D cases with curved membrane surface, the simulation details are similar, except that the binding probability needs to be integrated over the accessible area on the surface (for spheres we can use the closed-form expression Eq (S2)). We use a triangular mesh to build up the curved surface [43], thus the accessible area is composed of many triangular elements. Each element contributes to the binding probability by pair association probability *p_react_*(Δ*t*|*r_i_*) multiplying element area *s_i_* and the local density of lipids *ρ_i_*. Then the total binding probability is accumulated over all these elements based on Eq 6: 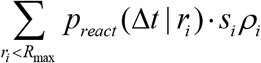. For curved membranes with analytical shapes such as the sphere, the reaction probability can be calculated analytically for each height *z*_0_ (Supporting Information).

For simulations of explicit protein particles restricted to the 2D surface with implicit lipids, simulations are similar to 3D cases, but *D_z_*=0 and distances are based only on the 2D separation between points. When protein particles bind to the lipids, their position does not need to change, but their diffusion coefficient is updated due to formation of a complex and they are no-longer free to bind. Dissociation then reverses these changes, without requiring specific placement of the protein.

### D. Simulation details of the Surface Adsorption model

For simulations of surface adsorption using the 1D GF of section IID, the binding probability of a solution particle is evaluated by Eq. 25. If the binding occurs, the particle will be put at the nearest point on the surface *z* = *σ* (i.e. radiation boundary condition). The position for this 1D approach after a survival would require a free diffusion move in *x* and *y*, and a movement in *z* sampled from the *p_irr_* solution shown in Eq. 24 for a new *z* value by solving 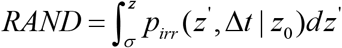, where *RAND* is a uniformly distributed random number between 0 and the value of the survival probability 1 − *P_AD-GF_* (Δ*t* | *z*_0_). This sampling ensures an exact solution to the model defined in Section D1, and is the same approach used in GFRD [23]. For the bound particle, the dissociation probability is evaluated by Eq. (3). If the dissociation occurs, the particle becomes free, but remains in contact with the surface at *z* = *σ*. In the next time-step it can then either react or diffuse.

### E1. Triangulated mesh model for complex surfaces

A subdivision limit surface (SLS) mesh algorithm described in [55, 56] was used for those complex surfaces not representable by a closed form. Briefly, the SLS is determined by subdividing each triangle of a mesh into four smaller triangles repeatedly. Each iteration of the subdivision creates a finer and finer mesh. A rule is applied to generate the vertices of the subdivided triangles from the previous iteration. That rule is codified as a matrix transformation of the previous iteration’s vector of vertex coordinates that yields the coordinates of the new vertices. Taking the limit of infinite applications of the rule yields a set of points constituting a surface that is tangent plane continuous. In practice, the limit surface itself is written as a polynomial spline that is proportional to the original vertices. The spline can be computed from the mesh that results following any amount of subdivision.

### E2. Detecting collisions with the triangulated mesh surface

At any iteration of subdivision, the spline of the limiting surface is contained completely within the convex hull defined by the spline control points (that is, the vertex coordinates of that subdivision). Therefore, if the convex hull of a second object does not intersect the convex hull of the control points, it does not intersect the limit surface (Fig. S4a). If an object does intersect the limit surface, it will intersect the convex hull of the mesh at any level of subdivision (Fig. S4b). Therefore, collisions of any convex object with the mesh can be excluded to arbitrary thresholds by checking for intersections between convex hulls at a particular depth of subdivision. Here the Gilbert-Johnson-Keerthi algorithm [44] was used to detect convex hull collisions. A recursive function was defined to detect collisions with the target object and the box spline of a mesh element. If a collision was detected at a particular subdivision level, the recursive function was called on each of the four subdivisions of the triangle. If collision is excluded the recursive branch terminates (Fig. S5). This recursion proceeds until a particular depth level (ten) at which point a collision was declared. This basic framework allows for computing the point of closest approach of a particle to the surface (by running bisection to determine the largest sphere that does not contact the surface) as well as detecting the point of intersection of a linear path through the surface (using that line segment for collision detection) (Fig. S6).

Convenient surface coordinates for a face of the SLS are its (nonorthogonal) spline parameters, here denoted *u* and *v*. These take the place of *x* and *y* that might be used on a planar surface. Where the Laplacian is appropriate for describing the time dependence of diffusion for *x* and *y* coordinates on a Cartesian plane, diffusion on the SLS must be propagated using the *Laplace-Beltrami operator* as done in [46]. This corrects for both the units and nonorthogonality of the spline coordinate.

## IV. Results

### A. Implicit lipid vs explicit lipid models for planar systems

The implicit lipid model always recovers the proper equilibrium, thanks to the use of the detailed balance expression in Eq. 10 to define the reflection surface RS (Fig 5). For simulations with 200 solution particles and a range of lipid densities and reaction-rate parameters, we further show that the kinetics are in excellent agreement with the results of explicit lipid simulations (Fig 5). In Fig 5b, we simulate a diffusion limited reaction (*k_a_*=173.59 nm^3^/μs, *k_b_*=2.09 s^−1^, *K*_D_=0.02 μM) at increasing lipid densities, illustrating how even with a low concentration of lipids (100/200^2^ nm^−2^), the kinetics are nearly the same. In Fig 5c, we simulate a rate-limited reaction (*k_a_*=0.166 nm^3^/μs, *k_b_*=1.00 s^−1^, *K*_D_ =10 μM). Because rate-limited reactions are relatively insensitive to diffusion and the spatial distributions of reactants, the close agreement between the models here is less surprising. Rate-limited reactions also produce similar kinetics in a non-spatial, ODE simulation of the same system (Fig. S2a). Diffusion to the surface does start to have an impact when the separation from the surface can reach 2*μ*m, but the change is relatively small compared to the substantial difference observed in diffusion-limited reactions (Fig. S2b). For diffusion-limited reactions, most collisions are reactive, and thus the assumption of the ODE that the lipids are well-mixed, compared to localized to the surface, produces much faster kinetics.

**Figure 5.**
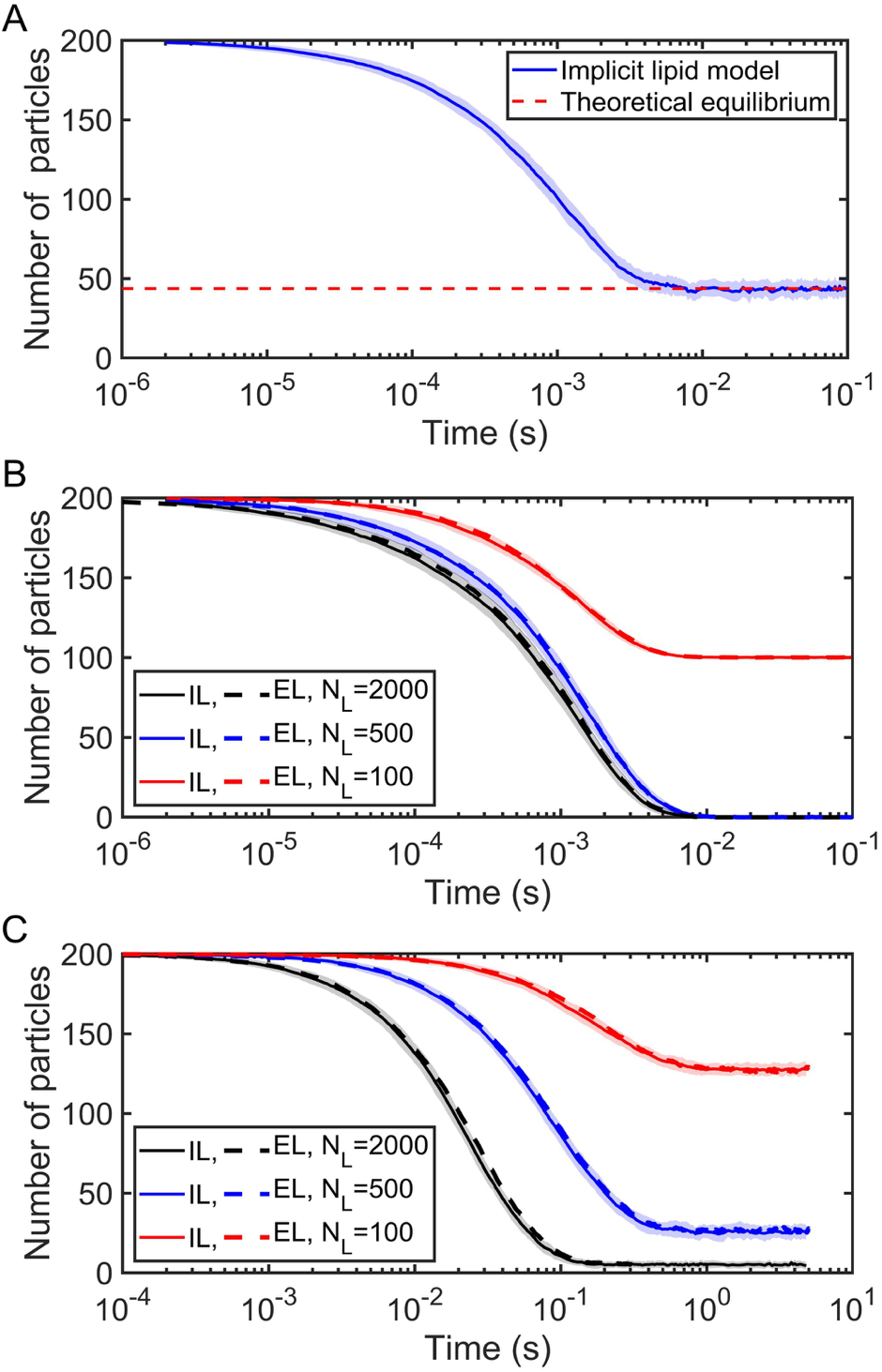
The implicit lipid (IL) and explicit lipid (EL) methods with the planar surface shown in Fig. 4A are in very close agreement for many-body simulations. The cubic system has a volume V=200^3^ nm^3^, surface area S=200^2^ nm^2^. All simulations use parameters: *σ*=1 nm, Δ*t* = 0.1 μs, 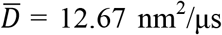 and particle number *N_A_* = 200 (all in solution at the beginning of simulations). (A) Kinetics of IL. Simulation parameters: *k_a_* =347.18 nm^3^/μs, *k_b_*=2.09×10^3^ s^−1^; the reflecting surface *RS*_3D_ = 0.69 nm; lipid number *N_L_*=500 (initially all unbound at the beginning of simulations). The equilibrium state is *K_D_* = *2k_b_/k_a_* = 20 μM, which is also defined as *K_D_* ≡ [A][L]/[AL] = *n_A_*[*N_L_*-(*N_A_*-*n_A_*)]/[*V*(*N_A_*-*n_A_*)], thus we have the number of free solution particles at equilibrium 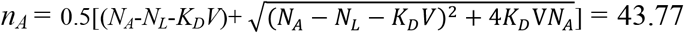. (B) Kinetics of IL and EL in diffusion-limited situations are also in excellent agreement: *k_a_* = 347.18 nm^3^/μs, *k_b_* = 2.09 s^−1^, *RS*_3D_ = 0.69 nm. (C) Kinetics of IL and EL in rate-limited situations: *k_a_* = 0.332 nm^3^/μs, *k_b_* = 1.00 s^−1^, *RS*_3D_ = 0.002 nm. In the figure, EL curves are shown as Mean calculated by 50 trajectories, while IL curves are shown as Mean ± Standard Deviation (SD) calculated also by 50 trajectories.

The CPU time of these simulations is independent of the number of lipids, and thus as the density of lipids or the size of the surface area increase, the implicit lipid model provides huge speed-ups relative to the explicit lipid model (Fig 6). The implicit model is actually slightly slower at the lowest lipid density; this is because most of the solution particles remain in solution, and must continue evaluating binding to the surface throughout the simulation, which requires more evaluations than if the proteins are bound to the surface. This is also why the explicit lipid simulations do not show linear slow-downs with increasing lipid copies—as more particles are bound to the surface, the speed of a time-step increases.

**Figure 6.**
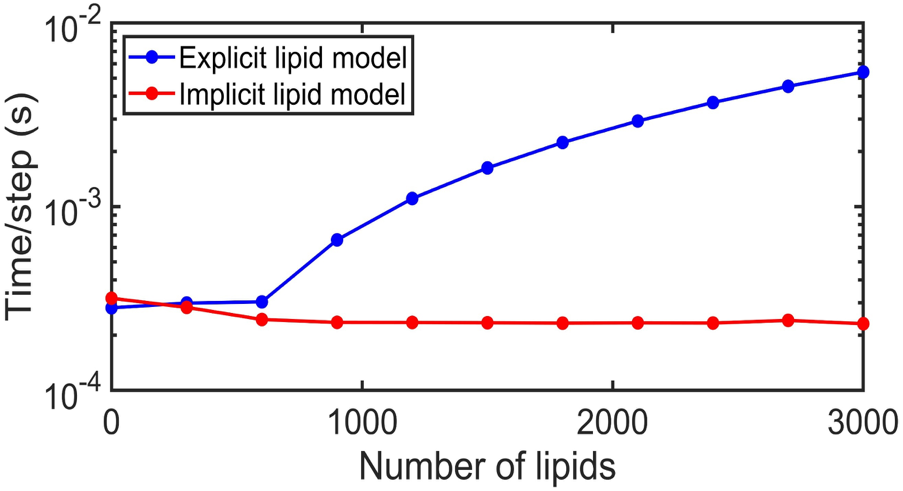
CPU time comparison for explicit lipid vs implicit lipid models. The implicit lipid model is independent of the number of lipids, whereas the explicit lipid model becomes slower with increasing lipids. Each simulation was run with 600 solution particles for 10^5^ steps in a cubic system (V=200^3^ nm^3^ and S=200^2^ nm^2^), with the average CPU time per step being plotted vs lipid copies.

### B. Adsorption method vs explicit lipid method for planar systems

For a standard surface adsorption model, the value of the adsorption coefficient *κ* does not change throughout the simulation [16]. This neglects occupancy of surface sites by other solution particles. For high lipid density, this does not matter and adsorption models produce very similar kinetics to the explicit lipid simulations, due to the excess of surface binding sites. However, when lipids are not in excess, the models diverge in both equilibrium and kinetics (Fig. 7). One adaptive approach to the adsorption model is to impose lipid density changes on it by dynamically setting *κ* = *k_on_*·*ρ*_L_(*t*) (The macroscopic binding rate *k_on_* stays the same as in the explicit lipid method), to account for occupancy of lipid binding sites.

**Figure 7.**
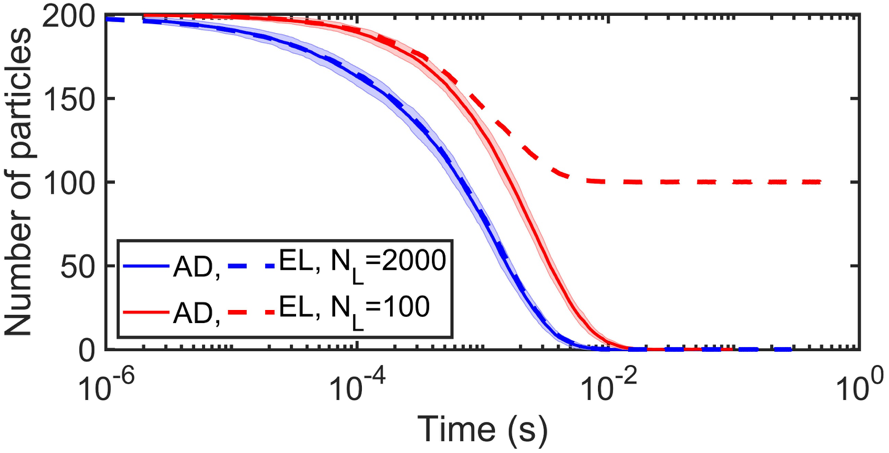
Explicit lipid (EL) method compared to surface adsorption (AD) model using Ref [16], where adsorption rate is fixed. For limited lipids, the fixed adsorption model produces a distinct equilibrium (red curve), because it does not account for occupancy of lipid binding sites. With high lipid density, the models converge (blue). All simulations use parameters: *σ* =1 nm, Δ*t* = 0.1 μs; cubic system with the volume V = 200^3^ nm^3^, surface area S = 200^2^ nm^2^; particle number *N_A_* = 200 (all in solution at the beginning of simulations). For explicit lipid model, we use 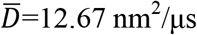 (diffusion constant of a solution particle with *D*=12 nm^2^/μs relative to a lipid on membrane with *D*=1 nm^2^/μs), *k_a_* = 347.18 nm^3^/μs, *k_b_* = 2.09 s^−1^, and thus the equilibrium *K_D_* =2*k_b_/k_a_* = 0.02 μM. But for adsorption model, we use *D*=12 nm^2^/μs (diffusion constant of a single particle in solution, not considering explicit lipids on membrane). For the limited lipids case, *ρ_L_* = = 0.0025 nm^−2^; for the sufficient lipids case, *ρ_L_* = 0.05 nm^−2^.

Using the surface adsorption model described in Section D1 (the 1D Smoluchowski model) with adaptive *κ* produces the same equilibria as the explicit lipid model for all lipid densities (Fig 8). The kinetics of the surface adsorption model are faster when the reaction is diffusion-limited (large *k_on_*) and lipids are low density (Fig 8a). As lipid density (and thus *κ*) are increased, the models converge. For rate-limited reactions (small *k_on_*) both models have similar kinetics even for low lipid densities, which is again not too surprising considering that these reactions are not highly sensitive to the specific particle distributions (Fig 8b). We note that *κ* is not unique to a single lipid density, and a lower density *ρ_L_* with stronger binding *k_on_* can produce the same *κ*. The models in Fig 8c have similar kinetics but distinct equilibria due to the use of adaptive *κ*, otherwise they would be identical. A major shortcoming of surface adsorption models is that they can only describe binding from a particle diffusing above the membrane and thus colliding with it. Proteins that are localized to the membrane surface are not freely diffusing in the direction normal to the surface, and for these molecules to bind to additional lipids, we can apply our implicit lipid model, below.

**Figure 8.**
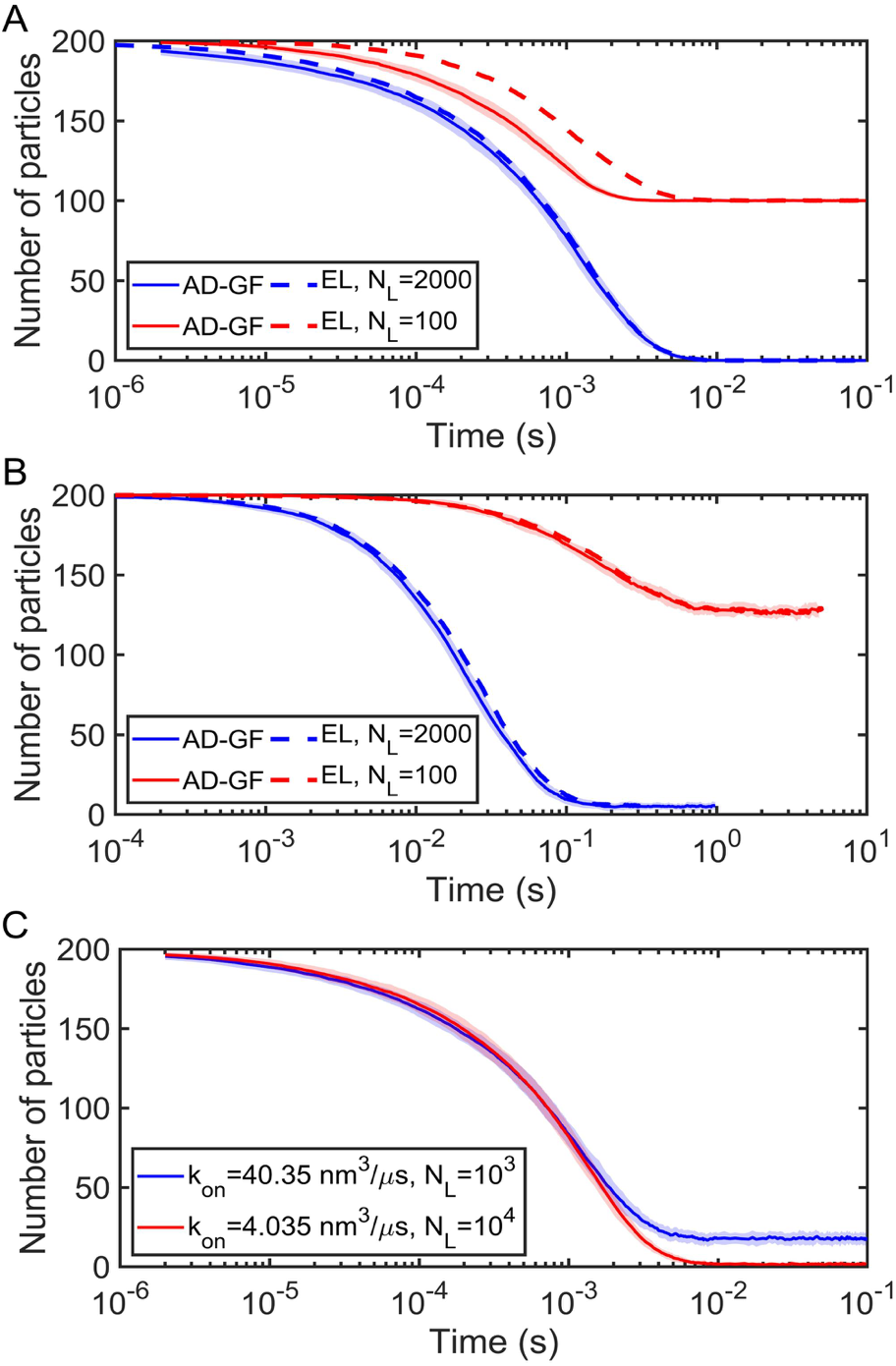
Surface adsorption model using the 1D Smoluchowski model (AD-GF) compared to explicit lipid (EL) method for planar surface system (Figure 4A). Adsorption rate is adaptive with lipid density. All simulations use parameters: *σ* = 1 nm, Δ*t* = 0.1 μs, and cubic system with volume V = 200^3^ nm^3^, surface area S = 200^2^ nm^2^, particle number *N_A_* = 200. For AD-GF, we use *D*= 12 nm^2^/μs, and for EL model 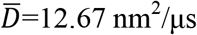. (A) Diffusion-limited situations. Parameters for AD-GF: *k_a_* = 173.59 nm^3^/μs, *k_b_* = 2.09 s^−1^, and the equilibrium *K_D_* =*k_b_/k_a_* = 0.02 μM. Parameters for EL, *k_a_* = 347.18 nm^3^/μs, *k_b_* = 2.09 s^−1^, and *K_D_* =2*k_b_/k_a_* = 0.02 μM. (B) Rate-limited situations. Parameters for AD-GF: *k_a_* = 0.166 nm^3^/μs, *k_b_* = 1 s^−1^, and the equilibrium *K_D_* =*k_b_/k_a_* = 10 μM. Parameters for EL, *k_a_* = 0.332 nm^3^/μs, *k_b_* = 1 s^−1^, and *K_D_* =2*k_b_/k_a_* = 10 μM. (C) Microscopic split of the adsorption coefficient *κ*(*t*) in AD-GF model. Blue curve used *k_a_* = 173.59 nm^3^/μs, *k_b_* = 2.09 s^−1^, *N_L_*=1000, *ρ_L_*=*N_L_/S*=0.025 nm^−2^, thus *k_on_*= 40.35 nm^3^/μs (evaluated by Eq. 21), which gives the adsorption coefficient *κ*=*k_on_·ρ_L_*=1.01 nm/μs. The red curve used *k_a_* = 8.53 nm^3^/μs, *k_b_* = 2.09 s^−1^, *N_L_*=10000, *ρ_L_*=*N_L_*/*S*=0.25 nm^−2^, thus *k_on_*= 4.035 nm^3^/μs, which also gives the adsorption coefficient *κ*=*k_on_*·*ρ_L_*=1.01 nm/μs at the beginning of the simulations. However, their kinetics and equilibrium states are different due to adapting to lipid occupancy. In the figure, each EL curve is shown as Mean calculated by 50 trajectories, while AD-GF curves are shown as Mean ± SD calculated also by 50 trajectories

### C. Binding to lipids in 2D: implicit vs explicit lipid model

In addition to binding from solution to a surface, we demonstrate that the implicit lipid model can also accurately reproduce the ability of particles, previously localized to the surface through one interface, to perform 2D binding reactions with the surface using another interface. In Fig. 9, we compare the results of implicit lipid model 2D with the explicit lipid method. For all the systems, whether diffusion-limited (*k_a_*=86.80 nm^2^/μs), rate-limited (*k_a_*=0.083 nm^2^/μs), or diffusion influenced (*k_a_*=8.68 nm^2^/μs), the implicit lipid model produces the proper equilibrium states due to enforcing detailed balance through the *RS*_2D_ parameter. For these 2D reactions, we find the kinetics of the implicit lipid model are slower than the explicit lipids, particularly for diffusionlimited reactions. This is likely due to the fact that in 2D, reaction dynamics are always sensitive to the spatial distribution of particles and are not characterized by a single macroscopic *k_on_* [53]. By replacing individual particles with a field, we eliminate rapid reactions due to pairs of particles that are close together, the effect of which is exaggerated in 2D relative to 3D, and by assigning *RS*_2D_ ≥ *σ* to ensure equilibrium, we also slow the reaction kinetics. *RS*_2D_ here increases from *σ* at the smallest *k_a_* value, up to 1.46*σ* for the largest *k_a_* (see Fig. S1).

**Figure 9.**
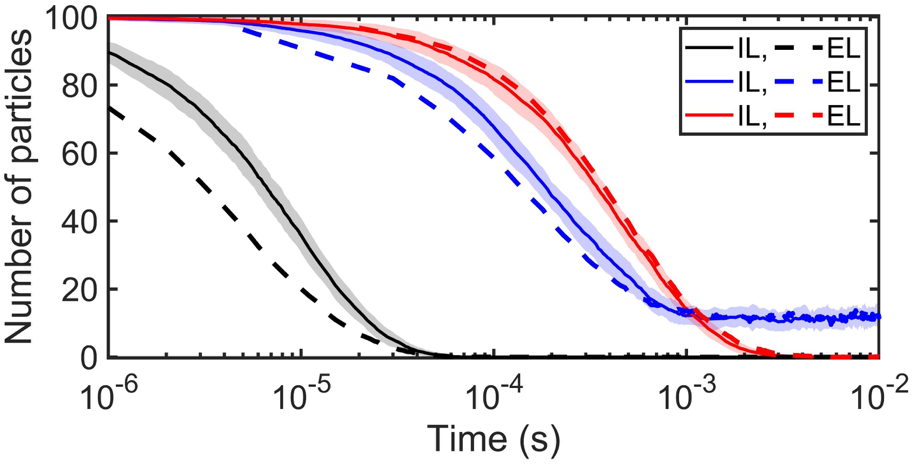
The 2D implicit lipid (IL) model reproduces the proper equilibrium as the explicit lipid (EL) model, but the kinetics are slower for diffusion-limited systems. The system used the flat surface as shown in figure 4A. All simulations used parameters: *σ* = 1 nm, *D* = 1 nm^2^/μs, Δ*t* = 0.1 μs, particle number *N_A_* = 200, lipids number *N_L_*=1000 (all are free at the beginning of simulations). Three different simulations are carried out. The strongly diffusion influenced black curve used *k_a_* = 86.80 nm^2^/μs, *k_b_* = 2.09 s^−1^, the equilibrium *K_D_* =*k_b_/k_a_* = 2.41 ×10^−2^ μm^−2^, surface area S= 200^2^ nm^2^ and the reflecting distance RS_2D_= 1.46 nm. The diffusion-influenced blue curve used *k_a_* = 8.68 nm^2^/μs, *k_b_* = 2.09×10^3^ s^−1^, *K_D_* =*k_b_/k_a_* = 2.41 ×10^2^ μm^−2^, S = 700^2^ nm^2^ and RS_2D_= 1.35 nm. The rate-limited red curve used *k_a_* = 0.083 nm^2^/μs, *k_b_* = 1 s^−1^, *K_D_* =*k_b_/k_a_* = 12.05 μm^−2^, S = 200^2^ nm^2^ and RS_2D_= 1.00 nm. In the figure, each EL curve is shown as Mean calculated by 50 trajectories, while IL curves are shown as Mean ± SD calculated also by 50 trajectories, and SD is shown in shaded region

### D. Implicit lipid vs explicit lipid model on curved surfaces

In Fig. 10, we again find excellent agreement between the implicit and explicit lipid simulations, but here with binding to curved surfaces (Fig. 4c-d). As we noted above, the implicit lipid reaction probabilities are analytical for a sphere but must be numerically integrated over the separation between the solution molecule and the surface for the triangulated mesh surface of Fig 4d. For the explicit lipid simulations, the lipids must also perform diffusion on a curved surface. The implicit lipid model produces exact equilibrium populations and excellent agreement with the kinetics of the explicit lipid method, whether in spherical systems (Fig. 10a-d), in a cylindrical tube (Fig S7), or with an irregular curved surface (Fig. 10e).

**Figure 10.**
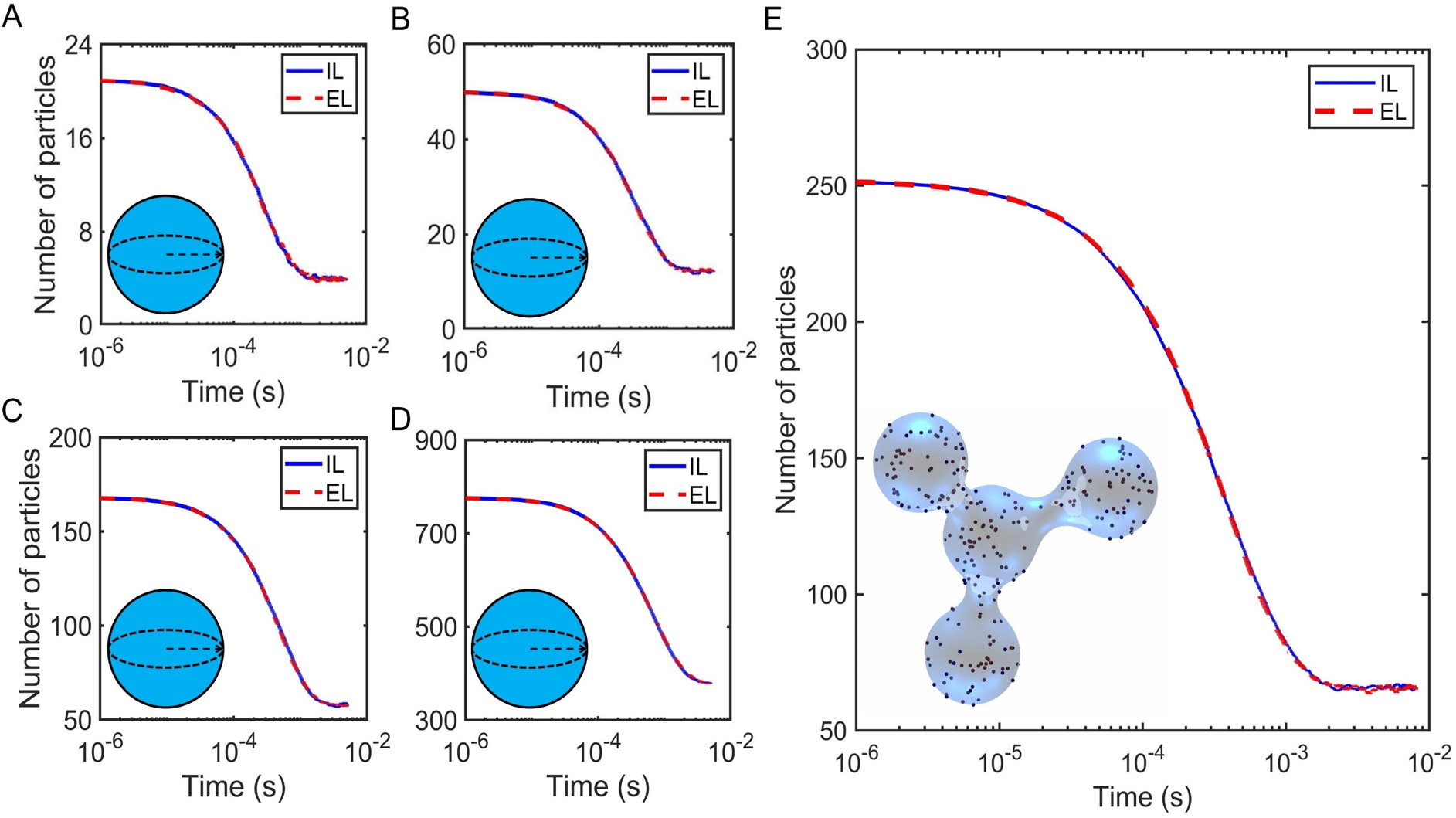
For binding to curved surfaces, the implicit lipid (IL) model shows very close agreement with the explicit lipid (EL) model results, with curved surfaces shown as figure 4C and 4D. All simulations used parameters: *σ* = 1 nm, 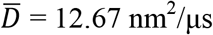, Δ*t* = 0.1 μs, *k_a_* = 347.18 nm^3^/μs, *k_b_* = 2.09×10^3^ s^−1^, thus the equilibria *K_D_* =2*k_b_/k_a_* = 20 μM. (A) Spherical surface with the radius R = 75 nm. The solution particle number *N_A_* = 21, lipids on surface *N_L_*=112. Particles diffuse inside the sphere or diffuse while bound onto the sphere surface, but cannot move outside the sphere. At the beginning of the simulations, all solution particles and lipids are unbound. (B) Spherical surface, R = 100 nm, *N_A_* = 50, *N_L_*=199. (C) Spherical surface, R = 150 nm, *N_A_* = 168, *N_L_*=448. (D) Spherical surface, R = 250 nm, *N_A_* = 776, *N_L_*=1244. (E) Irregular surface. Particles are closed inside the surface and diffuse within the enclosed volume or on while bound to the surface. The total volume accessible for solution particles, is V = 2.19× 10^7^ nm^3^ and total surface area S = 6.02×10^5^ nm^2^. Solution particle number *N_A_* = 252, lipids on surface *N_L_*=934. Each EL curve is the mean calculated by 100 trajectories, and each IL curve is the mean from 100 trajectories.

### E. Implicit lipid vs explicit lipid with non-uniform lipid distribution

Finally, in Fig. 11 we illustrate how the implicit lipid approximation also applies to surfaces with a non-uniform distribution of lipids, which could occur due to lipid phase separation or lipid microdomains. Here, the explicit lipid model had individual lipids restricted to one patch of the surface, and the implicit lipid model thus had the density also restricted to a patch of the surface (Fig 4b). This model is also an important demonstration of how this method can be coupled to a hybrid single-particle/PDE based approach, with lipid densities that can vary along the surface. The implicit lipid model shows excellent agreement with the explicit lipid method for both rate-limited and diffusion limited reactions.

**Figure 11.**
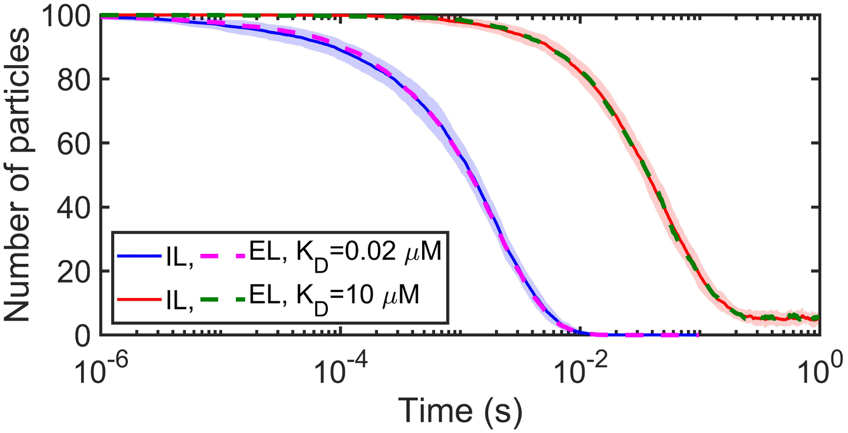
The implicit lipid (IL) model agrees very well with the explicit lipid (EL) method for an inhomogeneous lipid distribution generated by restricting lipids to half the surface, as shown in Figure 4B. The box system has a volume V=200^3^ nm^3^, surface area S = 200^2^/2 nm^2^. All simulations use parameters: *σ* = 1 nm, *D* = 12.67 nm^2^/μs, Δ*t* = 0.1 μs, solution particle number *N_A_* = 100, lipids on surface *N_L_*=1000. The blue and pink curves used *k_a_* = 347.18 nm^3^/μs, *k_b_* = 2.09 s^−1^, *K_D_* =2*k_b_/k_a_* = 0.02 μM, and RS_3D_=0.69 nm. The red and green curves used *k_a_* = 0.332 nm^3^/μs, *k_b_* = 1 s^−1^, *K_D_* =2*k_b_/k_a_* = 10 μM, and RS_3D_=0.002 nm. Each line is Mean ± SD from 50 trajectories.

## V. Discussion/Conclusions

Single-particle RD methods are a powerful tool for studying cell-scale dynamics, and by integrating them with increasingly advanced membrane models, we push the development of cell-scale membrane remodeling simulations. We here introduced an implicit lipid RD model for reversible binding of particles to membranes of varying lipid density and curvature, demonstrating that we could quite accurately reproduce the equilibrium and kinetics of explicit lipid simulations at speed increases of orders of magnitude. This method enables simulations of proteins interacting with large and arbitrarily shaped membranes without the considerable expense of propagating lipid binding sites. Unlike models of surface adsorption, we developed our method to integrate over all possible collisions with the surface even for large time-steps, thus ensuring and demonstrating its ability to apply to curved or inhomogeneous membranes, with excellent agreement to explicit lipid binding kinetics. Uniquely, our method also allows for binding between proteins on the surface and additional lipids (a 2D reaction), which represents an important stabilizing contact for proteins that are initially localized to the membrane via a protein binding partner [7]. Our RD code is open source and incorporated into a new software tool to facilitate user-defined models, and is immediately downloadable from Github at https://github.com/mjohn218/fpr_implicit, including both the implicit lipid model and the surface adsorption model for comparisons.

To ensure that our implicit lipid model here is transferrable and comparable to other rate-based tools, whether other single-particle [11, 23, 24, 33, 35], RDME [29–31], or ODE/PDE based tools [25, 26], and quantitatively comparable to experimental studies of binding [57], we have taken care to quantify how our model parameters translate to macroscopic rates and equilibrium constants. We additionally characterized how surface adsorption rates, which have distinct units relative to bimolecular association rates, can be adapted to, if necessary, better mimic the binding and kinetics of explicit lipid simulations. A limitation of our approach is that the implicit lipid model, unlike binding models between explicit particle pairs [58, 59], is not a formally exact solution to a well-defined dynamical system. The particles are propagated according to free diffusion, which is algorithmically much simpler and transferrable to other RD tools, but neglects the effect of the reactive lipids on particle positions. We correct for this approximation by introducing the *RS* distance in both 3D and 2D, to ensure detailed balance between binding and unbinding is maintained. Remarkably, this model nonetheless produces kinetics that are in very close agreement with explicit lipid models, indicating the robustness of replacing individual particle interactions with their spatially integrated average. An exception is for purely 2D reactions, where the implicit lipid model has slower kinetics for diffusion-limited reactions, likely due to the sensitivity of reaction dynamics in 2D to spatial distributions [53, 60].

A limitation of our theoretical approach is that for binding to curved surfaces, we approximate one aspect of the model, the pairwise reaction probability (Eq. 2), as unaffected by curvature. We do account for the effect of curvature on determining separation from the membrane and thus the integrated reaction probability (Eq 11), on placement of particles that displace beyond the membrane, and on the macroscopic rate (Eq 21). Correcting the pairwise reaction probabilities (Eq. 2) for curvature would be a numerically expensive calculation that would introduce minor changes. In our simulations of binding on curved surfaces, our results for both implicit and explicit-lipid models produced excellent agreement with the theoretically predicted equilibrium, indicating that these curvature effects are quite small, even for a sphere of R=75nm. Reducing the time-step reduces the scale of the local curvature, thus improving the assumption of a planar reflecting surface on pairwise reaction probabilities and providing a simple mechanism for evaluating the impact of curvature on binding. We found that using the reaction probabilities from the model of Eq 1 worked even for our largest time-step of 1 *μ*s, and were independent of time-step, as expected for a convergent method (Fig S8). We acknowledge that our algorithm for arbitrary curved surfaces does require numerical integration for determining reaction probabilities, which is more expensive than simple adsorption models, but is nonetheless straightforward to manage using widely existing libraries such as GSL. We also note that one advantage of surface adsorption methods is that they can more easily introduce forces between planar surfaces and solution particles [18], because they use solutions to the 1D GF rather than the 3D version we use here.

Our implicit lipid model can be combined with complex networks of protein-protein interactions to efficiently simulate a range of systems involving reversible localization to the membrane, such as models of clathrin-cage assembly [7, 35, 61, 62] and cell polarization[37], or experiments involving binding and oligomerization on membranes [63]. Integration with continuum models of surfaces, as we have done here, is especially critical for developing quantitative and dynamical models of membrane dynamics as they are driven by proteins [64–67]. These models are key for understanding processes from endocytosis, exocytosis, and virion formation at the smaller scale [66–68], to cell division at the larger scale. These current models lack single-particle resolution of the proteins localizing from solution to the membrane, as we do here, which is a necessary step in capturing more molecular-level dynamics over cellular lengthscales. Also, although we have focused here on biological problems involving membranes, our rate-based approach applies equally well to studying any type of reaction-dynamics involving surfaces, such as heterogeneous catalysis [69, 70]. Single-particle RD is more expensive than PDE or RDME simulations of reaction dynamics, but it is unique in providing structure to species that can be used to capture short-range spatial correlations and structure-resolved self-assembly [35]. While RD methods in general lack the physico-chemical details of Molecular Dynamics approaches[8, 9], single-particle RD at least has the capacity to build in deterministic forces between species [14, 36], and RD dramatically exceeds Molecular Dynamics in reaching cellular length and time-scales, with applications to equilibrium and non-equilibrium systems. Our method here provides an essential component for performing tractable simulations involving large networks of proteins that localize dynamically to membranes, to ultimately build predictive models for cell biology.

## Acknowledgements

Research reported in this publication was supported by a National Science Foundation CAREER Award to M.E.J., award No. 1753174. The research used the NSF XSEDE computational resources of SuperMic under XRAC MCB150059, and the MARCC supercomputer at JHU. Funding for AJS, RK and GT was provided by the intramural research program of the *Eunice Kennedy Shriver* National Institute of Child Health and Human Development of the NIH.

## APPENDIX A: Implicit Lipid Model on Curved Surfaces

Our implicit lipid model defines the binding probability of a solution particle to a membrane surface (Eq. 5). For the planar surface, the binding probability is analytically expressed by Eq. (6). For an arbitrarily curved surface, we have to use Eq. (11) to numerically calculate the binding probability. However, for the spherical surface, an analytical solution exists we derive here, both for negative curvature (binding from within the sphere) and positive curvature (binding from outside the sphere). Together, the two cases also provide the reflecting surface for curvature of arbitrary sign, provided that *σ* ≪ *R*, or equivalently, that *σ* |*c*_1_| < 1 and *σ* |*c*_2_| < 1 where *c*_1_ and *c*_2_ are the two principal curvatures of the surface [71].

### 1. Implicit lipid model: Binding probability evaluated to a sphere from within

Consider a solution particle diffusing inside a spherical surface of radius *R*, and let *r* be the distance between the particle to the sphere center, here *r* < *R* (Fig. 12a). One ring patch on the surface has the length 2*πR*·sin*θ*, and width *R*·*dθ*, where *θ* is the polar angle. Thus this ring area is 2*πR*^2^sinθ·*dθ*. Then the lipid number on this ring area is *ρ_L_*·2*πR*^2^sin*θ*·*dθ*, where *ρ_L_* is the lipid density. The distance of the solution particle to any lipid on this ring area is 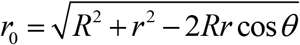, and their reacting probability is as the pair-wise one, *p_react_*(Δ*t*|*r*_0_) (Ref [15]) thus the binding probability of the solution particle to this ring area is accounted for all lipids on this ring area: *p_react_*(Δ*t*|*r*_0_) ·*ρ_L_*·2*πR*^2^sin*θ*·*dθ*. Then the total binding probability should be accumulated over the whole surface:

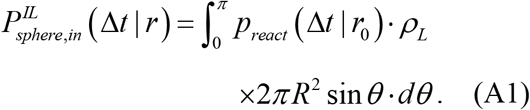

**Figure 12.**
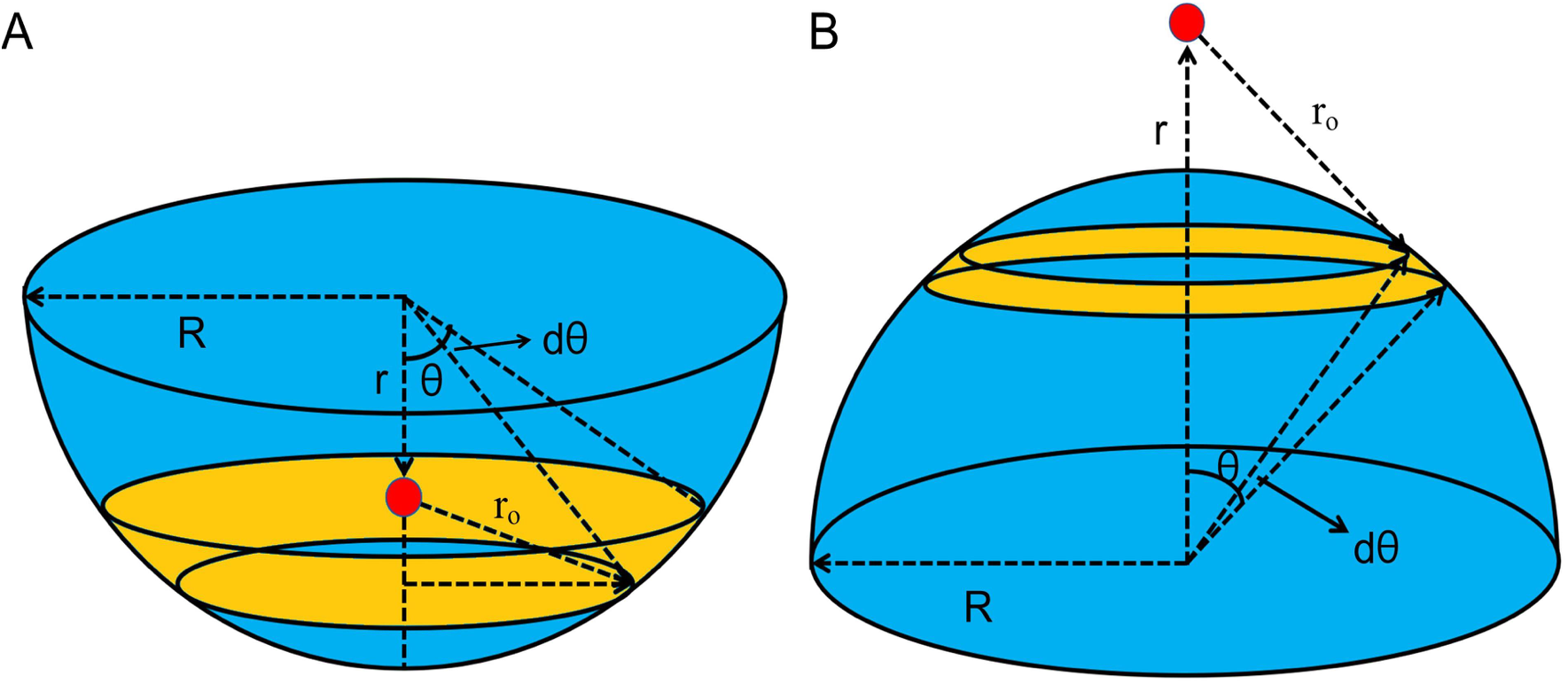
Geometric schematic of implicit lipid model with sphere surfaces. Binding to the surface from within (A) and from outside (B). The yellow represents one ring area on the surface, and the red point represents the solution particle.

Taking *p_react_*(Δ*t*|*r*_0_) [15] into Eq. (A1), we can have the analytical expression,

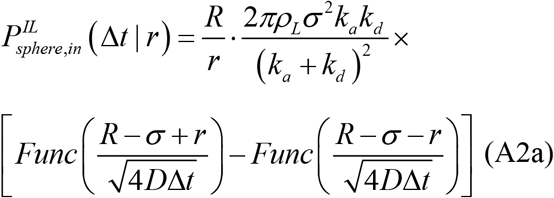

where *R* is the sphere radius, *r* is the distance between the solution particle and the sphere center, *ρ_L_* is the lipid density on the surface, *k_d_* =4*πσD*, and the function *Func* has the form of

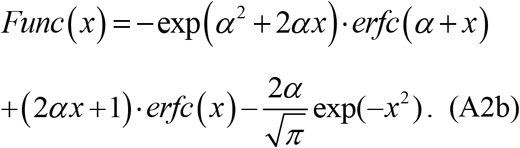

where 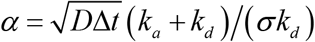. When the particle gets too close to the surface, *r* > *R* − *σ*, we set the binding probability is always as 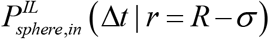. Importantly, if the sphere radius *R* → +∞, the sphere binding probability, Eq. (A2), near the surface is approximately equal to Eq. (8), the planar binding probability:

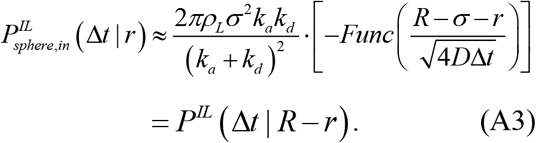

The reflecting surface *RS_sphere_* can be calculated according to the detailed balance. When the system is at equilibrium state, during the time Δ*t*, the quantity of particles binding to membrane should be equal to that of dissociation,

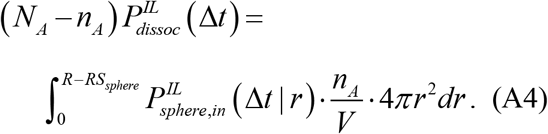

where the dissociation probability 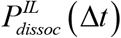 is described as Eq. (7), and V is the volume of system, *N*_A_ is the total number of solution particles, *n_A_* is the number of unbound particles at equilibrium state. The solution of *RS_sphere_* is

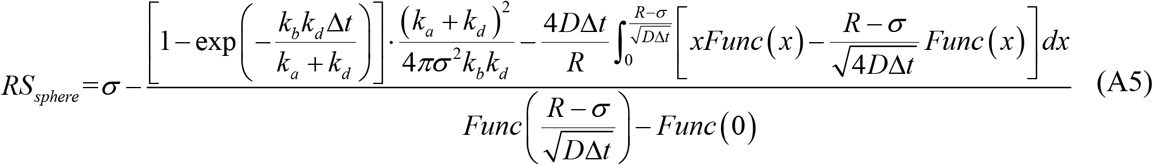

Generally, both *k_b_* and Δ*t* are small, and *R σ*, then *RS_sphere_* is approximately expressed as

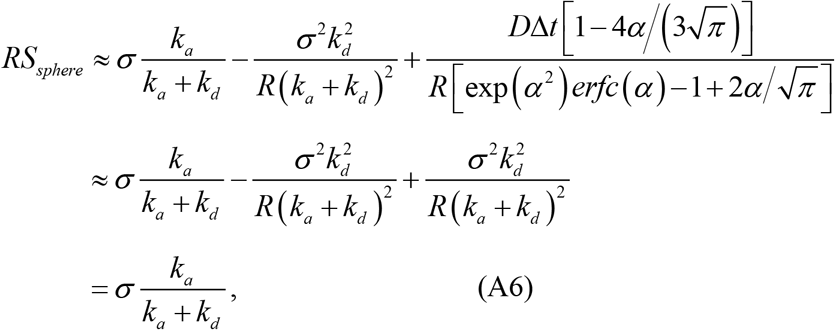

where *k_d_* =4*πσD*. As we can see, the expression of *RS_sphere_* is the same as the *RS*_3D_ for the flat surface situation. Note, to get the Eq. A6, we don’t have to use *R* →+∞, instead *R σ* is enough, thus the reflecting surface *RS* is not related to the membrane curvature.

### 2. Implicit lipid model: Binding probability evaluated to a sphere from outside

If we consider the situation that solution particles diffuse outside the spherical surface (Fig. 12b), the binding probability is calculated similarly as the one inside the sphere. One ring patch on the surface has the length 2*πR*·sin*θ*, and width *R*·*dθ* (*R* is the sphere radius, *θ* is the polar angle), thus the ring area is 2*πR*^2^sin*θ*·*dθ* and the lipid number on this ring area is *ρ_L_*·2*πR*^2^sin*θ*·*dθ*, where *ρ_L_* is the lipid density. The distance of the solution particle to any lipid on this ring area is 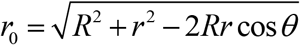 (*r* is the distance between the solution particle to the sphere center, here *r* > *R*), and their reacting probability is the same as the pair-wise one, *p_react_*(Δ*t*|*r*_0_). The binding probability of the solution particle to this ring area is: *p_react_*(Δ*t*|*r*_0_)· *ρ_L_*·2*πR*^2^sin*θ*·*dθ*. The total binding probability is integrated over the whole surface. Then the total binding probability is

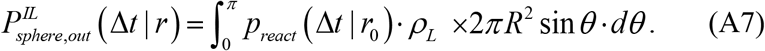

Taking *p_react_*(Δ*t*|*r*_0_) [15] into the above equation yields the analytical expression

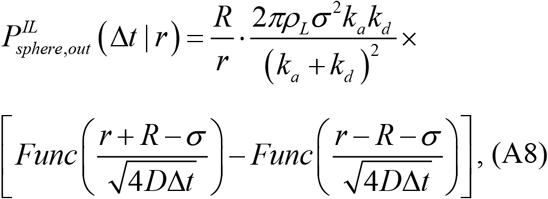

where *Func* is defined as in Eq. A2b. When the particle gets too close to the surface, *r* < *R* + *σ*, we set the binding probability to 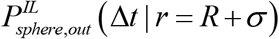. If the sphere radius *R*→+∞, and the solution particle is near the surface *r*-*R* ≪ *R*, then the binding probability is approximately equal to Eq. (8), the planar binding probability:

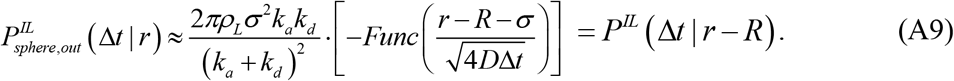

The reflecting surface is calculated the same as above, and when *k_b_* and Δ*t* are small, and *R σ*, we still have

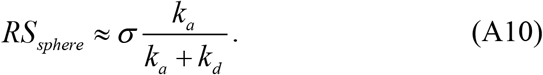

To sum up, when the fluctuation of the membrane surface is limited, and the local curvature is small, the probability of the solution particle binding to the curved surface, in our implicit lipid model, is similar to the particle binding to a flat surface, and the position of reflecting surface is also the same as the flat surface case.

