## Supplemental Figures S1-S8 for "An implicit lipid model for efficient reaction-diffusion simulations of protein binding to surfaces of arbitrary topology"

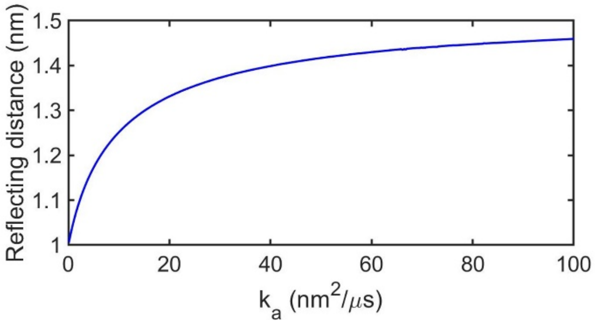


**Figure S1.** Reflecting distance $RS_{2D}$ of the 2D implicit lipid model, which must be $\geq \sigma$. Parameters: $\sigma$ = 1 nm, $D$ = 1 $nm^2/\mu s$, $\Delta t$ = 0.1 μs, $k_b$ = 2.09 $s^{-1}$, $\rho_L$ = 0.025 $nm^{-2}$ (lipids number $N_L$=1000, surface area S =$200^2$ $nm^2$).

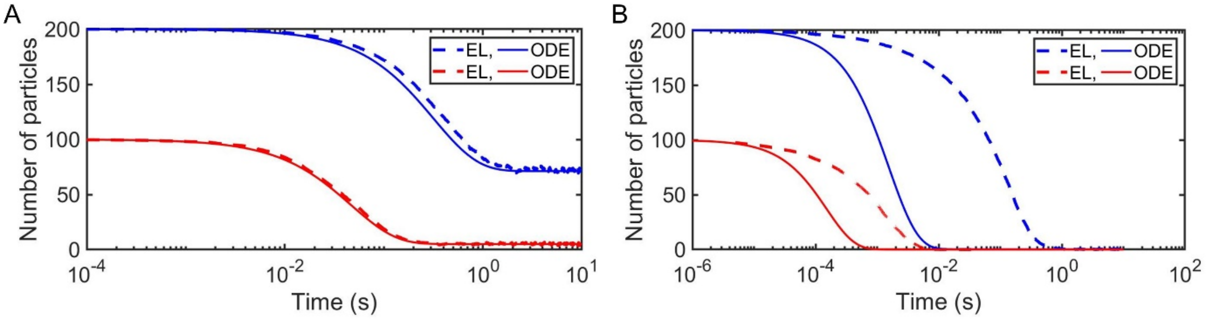


**Figure S2:** Kinetics of the explicit lipid (EL) model vs ODE simulations (solid lines). (A) Rate-limited systems show close agreement, where in both (A) and (B), red curves use a smaller cubic system V=$200^3$ $nm^3$, surface area S =$200^2$ $nm^2$, and blue curves use an elongated box with the z-dimension stretched to 2000 nm. ODE use $k_{on}$ =0.166 $nm^3/\mu s$, $k_b$=1.00 $s^{-1}$, $K_D = k_b/ k_a^{3D}$ =10 μM; EL use $k_a$ =0.332 $nm^3/\mu s$, $k_b$=1.00 $s^{-1}$, $K_D = 2k_b/ k_a$ =10 μM. (B) For diffusion-limited systems, the well-mixed ODE kinetics are much faster, as the EL model has to wait for solution particles to diffuse to the surface before productive reactive collisions with lipids can occur. ODE use $k_{on}$=54.58 $nm^3/\mu s$, $k_{off}$=0.657 $s^{-1}$, $K_D$ =0.02 μM; EL use $k_a$ =347.18 $nm^3/\mu s$, $k_b$=2.09 $s^{-1}$, $K_D$ =0.02 μM. All curves are shown as mean calculated by 50 trajectories. Parameters: $\sigma$ = 1 nm, $\overline{D}$= 12.67 $nm^2/\mu s$, $\Delta t$ = 0.1 μs, lipids number $N_L$=1000.

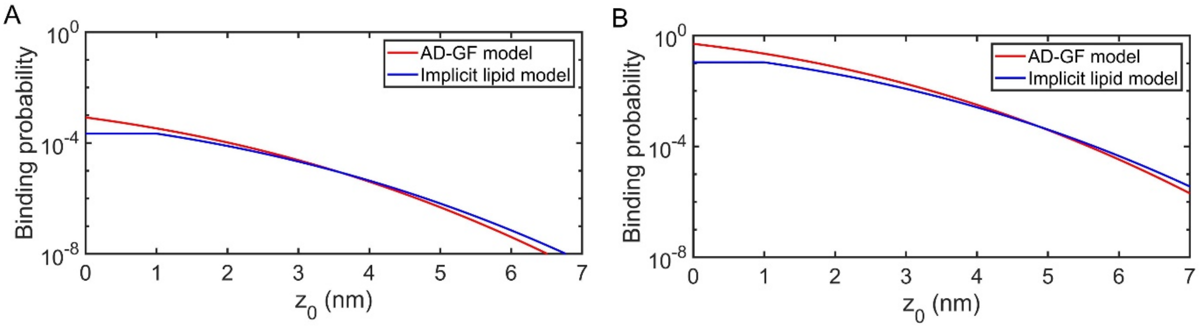


**Figure S3:** Binding probability of adsorption method using the 1D Smoluchowski model (AD-GF) compared to the implicit lipid model. Diffusion constants are $D_x$=$D_y$=$D_z$=12 nm$^2$/μs for the solution particle, and $D_x$=$D_y$=1 nm$^2$/μs, $D_z$=0 for the membrane lipids. This yields $D_{tot}$=12.67 nm$^2$/μs for the implicit lipid model, and $D_{tot}$=12 nm$^2$/μs for the AD-GF method. The time step is $\Delta t$ = 0.1 μs, and the lipid density is $\rho_L$ = 0.025 nm$^{-2}$. For the implicit lipid model $\sigma$ = 1 nm. For the AD-GF model we set $\sigma$ = 0 nm. (A) Rate-limited system. For AD-GF, $k_a$ = 0.166 nm$^3$/μs. For the implicit lipid model, $k_a$ = 0.332 nm$^3$/μs. (B) Diffusion-limited system. For AD-GF, $k_a$ = 173.59 nm$^3$/μs, and for the implicit lipid model, $k_a$ = 347.18 nm$^3$/μs.

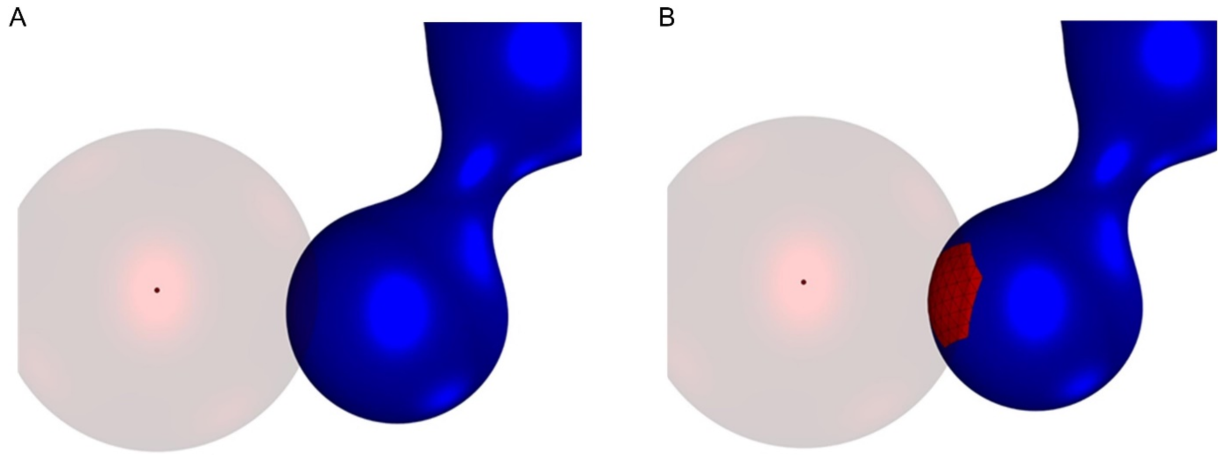


**Figure S4.** Computing the collision of a sphere with a surface is a basic element of preventing surface crossing as well as assembling the elements of a surface within some radius. To clarify the illustration, we show here the particle on the exterior of the blue membrane surface, although simulations were performed with particles inside the surface. (A) A radius does not intersect any of the convex hull of the target surface. This rigorously excludes any part of the surface being within that radius. (B) The sphere intersects the convex hull of at least one face. A solution particle is at the center of a translucent sphere. The algorithm tests for the intersection of the sphere and the convex hull of the surface faces at arbitrary recursion depth. The limit surface is shown in blue. Computation of the limit surface does not require recursion; it is an analytical form as a function of a coarse mesh.

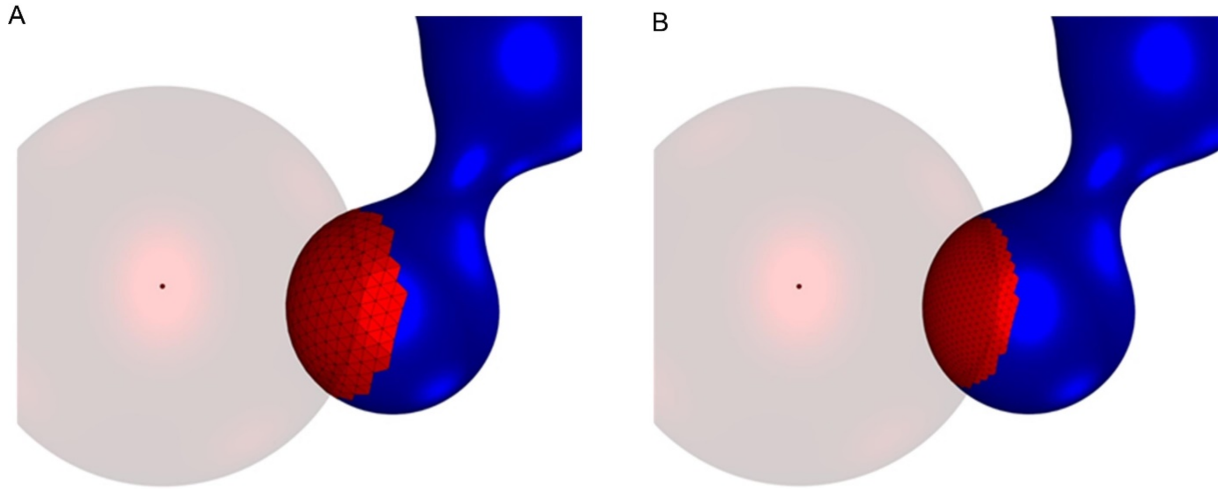


**Figure S5.** The recursion depth of the overlap search determines the accuracy of numerical integration of the binding probability. At left are the zero subdivision faces. At right a single subdivision yields four times as many smaller faces.

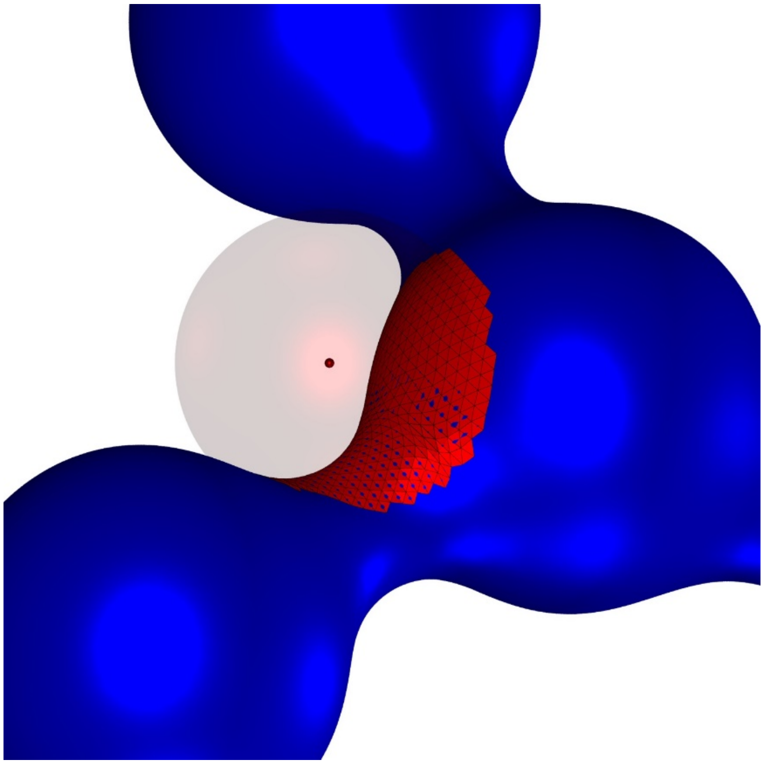

**Figure S6.** The algorithm can assemble a list of nearby faces for arbitrary shapes. In this image the blue limit surface overlaps the coarse triangular element at some faces yielding small blue patches. However, the entire surface is contained within the convex hull of the faces at any depth of recursion.

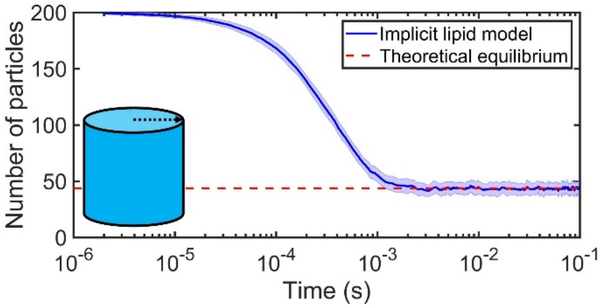


**Figure S7.** Simulation of implicit lipid model on a curved surface of cylinder. Simulation parameters: $\sigma$ = 1 nm, $D$ = 12.67 nm$^2$/μs, $\Delta t$ = 0.1 μs, and cylinder length L=200 nm, radius R=112.84 nm$^3$; solution particle number $N_{\mathrm{A}}$ = 200 and lipids number $N_L$=500 (all are unbound at the beginning of simulations); $k_a$ = 347.18 nm$^3$/μs, $k_b$ = 2.09×10$^3$ s$^{-1}$, thus the equilibrium state is $K_D = 2k_b/k_a$= 20 μM, which gives the number of free solution particles at equilibrium $n_{\mathrm{A}}\approx$ 43.77.

The curve is shown as Mean ± Standard Deviation (SD) calculated by 50 trajectories, and SD is shown in shaded region.

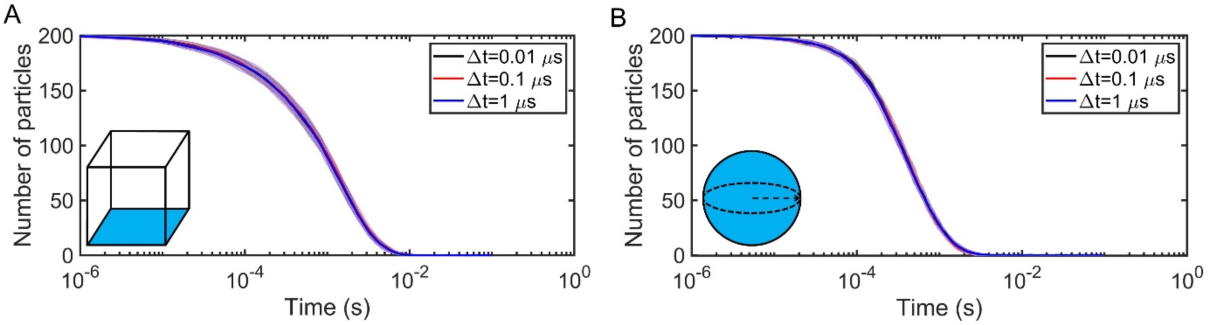


**Figure S8.** The size of the time-step does not influence kinetics of protein particles binding to membrane surface in the implicit lipid model. Parameters: $\sigma = 1$ nm, $D = 12.67$ nm$^2$/μs, $k_a$=347.18 nm$^3$/μs, $k_b$ = 2.09 s$^{-1}$, protein particle number $N_A$=200, lipids number $N_L$=500 (all particles are unbound at the beginning of simulations). For the cubic system, the volume V=200$^3$ nm$^3$, surface area S =200$^2$ nm$^2$; for the spherical system, the sphere radius R=124.07 nm. All curves are shown as Mean ±SD calculated by 50 trajectories, and SD is shown in shaded region.
